# Brain2GAN: Feature-disentangled neural encoding and decoding of visual perception in the primate brain

**DOI:** 10.1101/2023.04.26.537962

**Authors:** Thirza Dado, Paolo Papale, Antonio Lozano, Lynn Le, Feng Wang, Marcel van Gerven, Pieter Roelfsema, Yağmur Güçlütürk, Umut Güçlü

## Abstract

A challenging goal of neural coding is to characterize the neural representations underlying visual perception. To this end, multi-unit activity (MUA) of macaque visual cortex was recorded in a passive fixation task upon presentation of faces and natural images. We analyzed the relationship between MUA and latent representations of state-of-the-art deep generative models, including the conventional and feature-disentangled representations of generative adversarial networks (GANs) (i.e., *z*- and *w*-latents of StyleGAN, respectively) and language-contrastive representations of latent diffusion networks (i.e., CLIP-latents of Stable Diffusion). A mass univariate neural encoding analysis of the latent representations showed that feature-disentangled *w* representations outperform both *z* and CLIP representations in explaining neural responses. Further, *w*-latent features were found to be positioned at the higher end of the complexity gradient which indicates that they capture visual information relevant to high-level neural activity. Subsequently, a multivariate neural decoding analysis of the feature-disentangled representations resulted in state-of-the-art spatiotemporal reconstructions of visual perception. Taken together, our results not only highlight the important role of feature-disentanglement in shaping high-level neural representations underlying visual perception but also serve as an important benchmark for the future of neural coding.

**Author summary:** Neural coding seeks to understand how the brain represents the world by modeling the relationship between stimuli and internal neural representations thereof. This field focuses on predicting brain responses to stimuli (neural encoding) and deciphering information about stimuli from brain activity (neural decoding). Recent advances in generative adversarial networks (GANs; a type of machine learning model) have enabled the creation of photorealistic images. Like the brain, GANs also have internal representations of the images they create, referred to as “latents”. More recently, a new type of feature-disentangled “*w*-latent” of GANs has been developed that more effectively separates different image features (e.g., color; shape; texture). In our study, we presented such GAN-generated pictures to a macaque with cortical implants and found that the underlying *w*-latents were accurate predictors of high-level brain activity. We then used these *w*-latents to reconstruct the perceived images with high fidelity. The remarkable similarities between our predictions and the actual targets indicate alignment in how *w*-latents and neural representations represent the same stimulus, even though GANs have never been optimized on neural data. This implies a general principle of shared encoding of visual phenomena, emphasizing the importance of feature disentanglement in deeper visual areas.

## 1 Introduction

The brain is adept at recognizing a virtually unlimited variety of different visual inputs depicting different faces, objects and scenes, with each stimulus generating a unique pattern of neural activity. However, the complexity of multi-layered visual processing, occurring between stimulus and neural response, has hindered a comprehensive understanding of the transformation between the two. In the field of neural coding, our focus is to characterize the stimulus-response relationship that underlies the brain’s ability to recognize the statistical invariances of structured yet complex naturalistic environments. *Neural encoding* seeks to find how properties of external phenomena are processed in the brain [1, 2, 3, 4, 5, 6, 7, 8, 9, 10, 11, 12, 13, 14], and vice versa, *neural decoding* aims to find what information about the original stimulus is present in and can be retrieved from the recorded brain activity by classification [15, 16, 17, 18, 19], identification [20, 21, 22, 23] or reconstruction [24, 25, 26, 27, 28, 29, 30, 31, 32, 33, 34, 35, 36]. In classification, brain activity is taken to predict the category to which the original stimulus belongs, based on a predefined set of categories. In identification, brain activity is utilized to identify the most probable stimulus from a given set of available stimuli. In reconstruction, a literal replica of the original stimulus is recreated which involves the extraction of *specific stimulus characteristics* from neural data. Note that the latter problem is considerably harder as its solution exists in an infinitely large set of possibilities whereas those of classification and identification can be selected from a finite set. In both neural encoding and -decoding, it is common to factorize the direct transformation into two by invoking an in-between feature space (Figure 2). The rationale behind this is twofold:

1. *Efficiency*: modeling the *direct* stimulus-response relationship from scratch requires large amounts of training data (up to the order of millions) which is challenging because neural data is scarce. To work around the problem of data scarcity, we can leverage the knowledge of computational models (typically, deep neural networks that are pretrained on huge datasets) by extracting their feature activations to images and then aligning these with the elicited neural activity to those images during neuroimaging experiments, based on the systematic correspondence between the two. This correspondence is discussed under ’earlier work’.
2. *Interpretability*: the computational model whose features align best with neural activity can be informative about what drives the neural processing of the same stimulus (i.e., a data-driven approach). As such, alternative hypotheses can be tested about what drives neural representations themselves (e.g., alternative objective functions and training paradigms). This explanatory property can be limited when models are directly optimized on neural data (i.e., an exploratory approach) due to the complexity of the learned transformations.

The main aim of this study is to characterize high-level neural representations underlying perception, for which we analyzed the relationship between brain responses and various feature representations of recent generative models with different properties such as feature disentanglement and language regularisation, each of which captured a specific set of features and patterns about the visual stimuli. The representation that best predicted neural activity, by taking a linear combination of its features, was used to reconstruct perceived stimuli with state-of-the-art quality (Figure 1).

**Figure 1:**
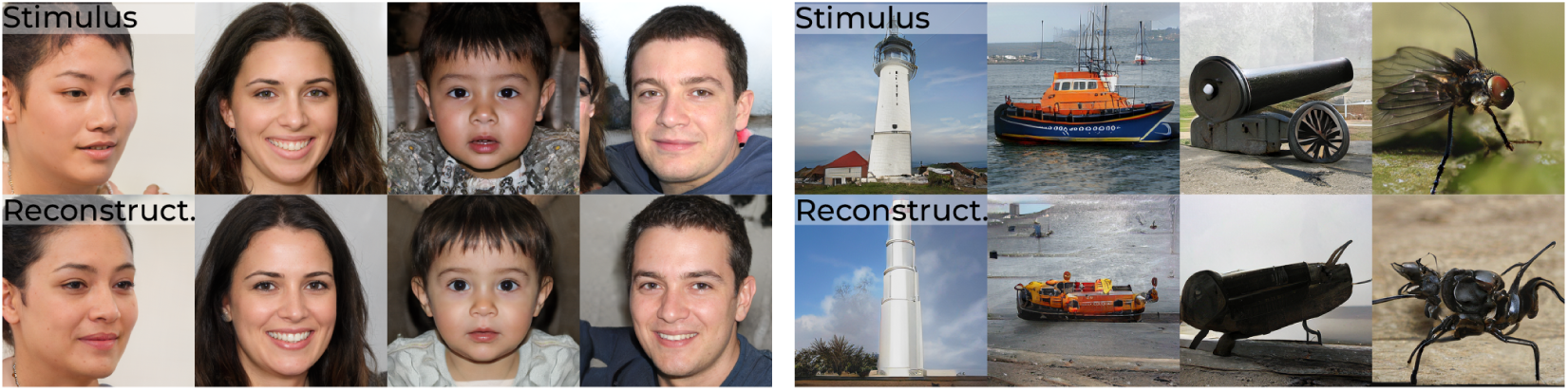
Example results. Stimulus (top) and reconstructions (bottom) from brain activity in V1, V4 and IT.

### 1.1 Modeling neural activity via feature-disentangled generative latents

Although neural representations are constructed from experience, an infinite amount of visual phenomena can be represented by the brain to successfully interact with the environment. That is, novel yet plausible situations that respect the regularities of the natural environment can also be mentally simulated or *imagined* [37]. From a machine learning perspective, generative models achieve the same objective by capturing the probability density underlying a huge set of observations. We can sample from this modeled distribution and synthesize new instances that appear as if they belong to the real data distribution yet are suitably different from the observed instances thereof. Particularly, generative adversarial networks (GANs) [38] are among the most impressive generative models to date which can synthesize novel yet realistic-looking images (e.g., images of human faces, bedrooms, cars and cats [39, 40, 41, 42] from latent vectors. In the context of generative models, like GANs, a *latent space* refers to a lower-dimensional data distribution (e.g., a standard Gaussian distribution) in which a more complex data distribution (e.g., face- or natural images) is encoded; it is a compressed and abstract space that captures the most essential features of the more complex data. A GAN consists of two neural networks: a generator network that synthesizes images from randomly-sampled latent vectors and a discriminator network that distinguishes synthesized from real images. During training, these networks are pitted against each other until the generated data are indistinguishable from the real data. The one-to-one (bijective) mapping from latents to images by the generator effectively models the ‘synthesis‘ operation (as specified in Figure 2) which can be exploited in neural coding to disambiguate the images from brain activity via their latents, since the visual content is deterministically specified by their underlying latents (such an approach was earlier suggested by [43]), and perform *analysis by synthesis* [44]. Note that, while the generator’s latent-to-image transformation performs the reconstruction of the perceived stimuli, it is the feature-response correspondence that enables the interpretation of neural activity as variations in the latent features.

Traditional GANs are known to suffer from *feature entanglement* where the generator has learned to fuse multiple features into a single latent dimension (i.e., a hyperplane in the multidimensional latent space) [45]. As a consequence of this fusion, the latent space contains biases inherited from the training dataset. To illustrate this, consider an example of generating images of human faces. A conventional GAN may entangle features like “gender” and “hair length” when predominantly exposed to feminine-looking faces with long hair and masculine-looking faces with short hair. Entanglement of these two features would result in biased outputs, hindering the generator’s ability to synthesize a masculine face with long hair, even if such combinations exist in reality. The concept of *feature disentanglement*, on the other hand, refers to the independence of different visual features, allowing variations in one feature to be untangled from others [46]. In a feature-disentangled GAN, the generator has learned to encode each facial feature independently. For example, changing the latent dimension corresponding to “hair length” would only modify the hair region of the generated face while keeping other features invariant. Here, we posit that feature-disentangled GAN latents exhibit a stronger alignment with neural representations in the ventral visual stream.

One member of the family of feature-disentangled GANs is StyleGAN [41] (Figure 3) - which maps the conventional *z*-latent via a multilayer perceptron (MLP) to an intermediate and less entangled *w*-latent space. Feature disentanglement is an emergent property that arose as the MLP learned to control diverse aspects of the image synthesis process within the training framework of StyleGAN. That is, the interplay between the generator’s evolving architecture, the injection of w-latents at different levels, and the network’s optimization for image generation contributes to the disentanglement of features in the w-latent space. Here, we propose feature-disentangled *w*-latents as a promising feature candidate to explain neural responses during visual perception. In brief, visual stimuli were synthesized by a feature-disentangled GAN and presented during a passive fixation task to a macaque with cortical implants in visual areas V1, V4 and IT (Figure 4). In contrast to many previous studies that relied on noninvasive fMRI signals with limited temporal resolution and low signal-to-noise ratio, the current use of multi-unit activity (MUA) [47] via 15 chronically implanted multielectrode arrays (each with 64 channels) provided opportunities for spatiotemporal analysis of brain activity in unprecedented detail. The electrode placings across these three visual areas are visualized in Figure 2. For neural encoding, we predicted brain activity from StyleGAN’s *z*- and *w*-latent representations, as well as Contrastive Language-Image Pre-training (CLIP; ViT-L/14@336px) latents which represent images and text in a shared representational space that captures their semantic relationships [48]. CLIP-latents are not just abstract representations of visual content, but they are also pivotal in the generative processes of contemporary latent diffusion models like Stable Diffusion [49]. Their key strength for our purposes lies in their ability to capture the essence of images in a way that reflects how the visual system of the brain processes visual inputs into semantic representations [50, 51].

**Figure 2:**
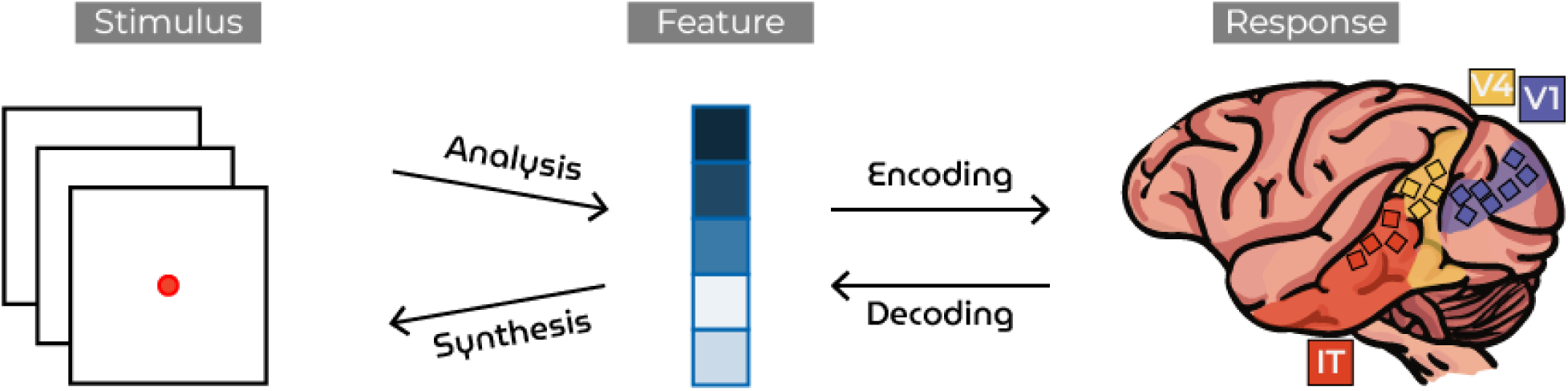
Neural coding. The transformation between sensory stimuli and brain responses via an intermediate feature space. Neural encoding is factorized into a nonlinear “analysis” and a linear “encoding” mapping. Neural decoding is factorized into a linear “decoding” and a nonlinear “synthesis” mapping.

**Figure 3:**
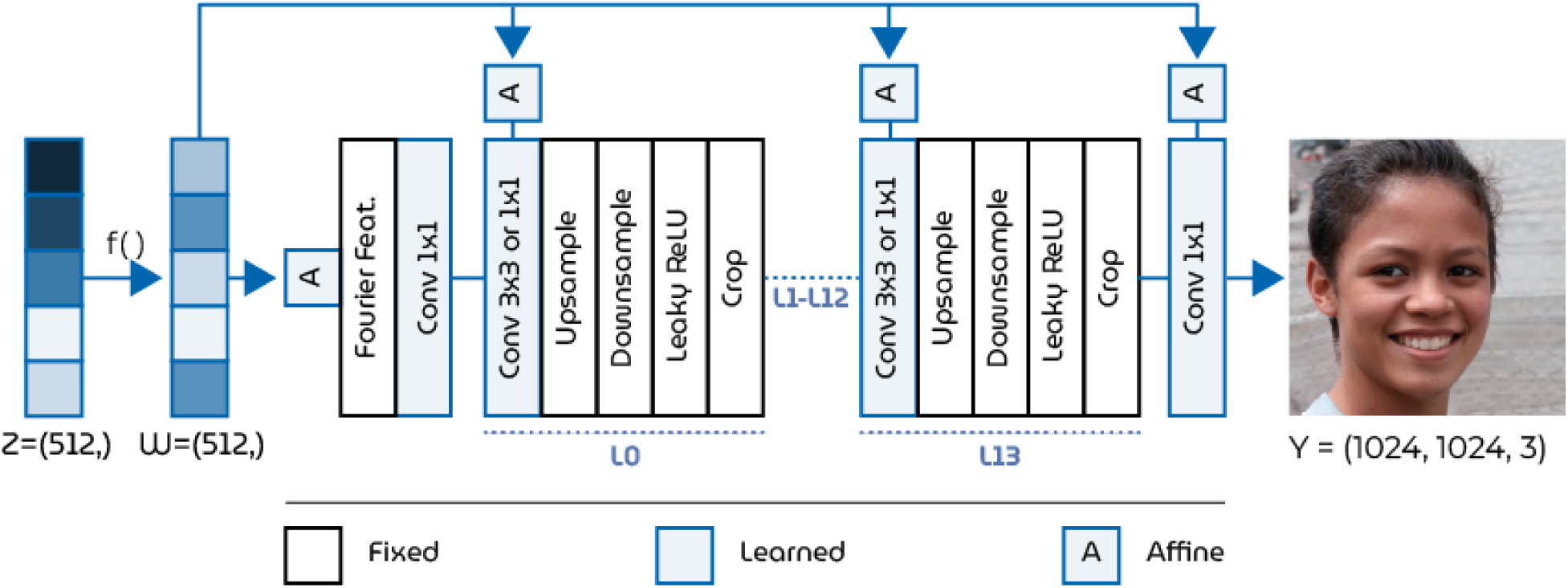
StyleGAN3 generator architecture. The generator takes a 512-dim. *z*-latent (entangled or correlated dimensions) as input and maps this to its 512-dim. *w*-latent (disentangled or decorrelated dimensions) via the MLP, *f* (), for feature disentanglement. Then, the *w*-latent is transformed into a 1024 *×* 1024 px RGB image.

**Figure 4:**
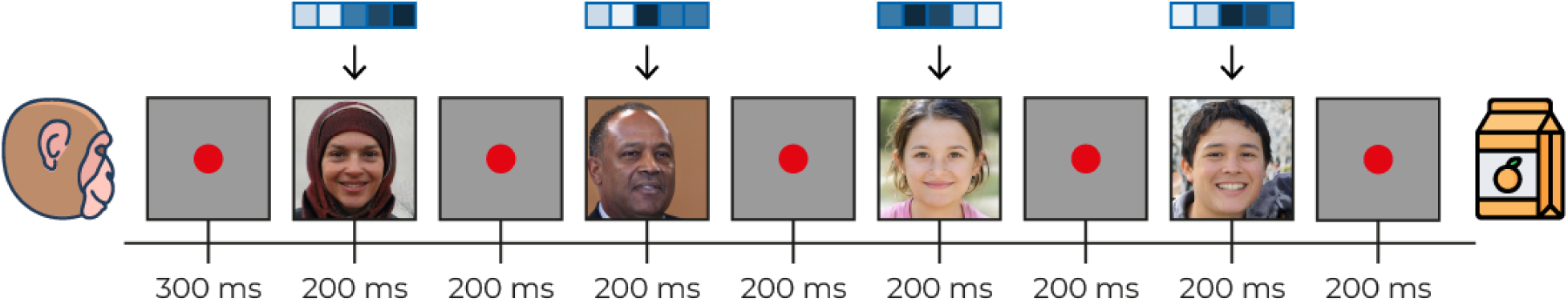
Passive fixation task. The monkey was fixating a red dot with gray background for 300 ms followed by a fast sequence of four face images (500^2^ pixels): 200 ms stimulus presentation and 200 ms inter-trial interval. The stimuli were slightly shifted to the lower right such that the fovea corresponded with pixel (150, 150). The monkey was rewarded with juice if fixation was kept for the whole sequence.

The contributions of this work are as follows: first, our encoding analysis revealed that *w*-latents, compared to the *z*- and CLIP-latents, were the most successful at predicting high-level brain activity in the inferior temporal (IT) cortex, which is located at the end of the visual ventral pathway. Second, neural decoding using *w*-latents resulted in highly accurate reconstructions that matched the stimuli in their specific visual characteristics. This was done by fitting a decoder to the recorded brain responses and the ground-truth *w*-latents of the training stimuli. We then used this decoder to predict the *w*-latents from responses of the held-out test set and fed these to the generator of the GAN for reconstruction [36]. Our findings indicate that the brain’s representation of visual information, in the context we studied, exhibits a degree of structured organization that aligns with our model, offering a new way forward for the previously limited yet biologically more plausible unsupervised models of brain function. Third, time-based neural decoding showed how the brain captured meaningful information about the stimulus in time. Finally, the interpretation of neural activity via the established response-latent relationship was explored by the application of linear operations to control specific visual features in the images. Taken together, the high quality of the neural recordings and feature representations resulted in novel experimental findings that not only demonstrate how advances in machine learning extend to neuroscience but also serve as an important benchmark for future research.

### 1.2 Earlier work

Visual experience is partially determined by the selective responses of neuronal populations along the visual ventral “what” pathway [52] where the receptive fields of neurons in early cortical regions are selective for simple features (e.g., local edge orientations [53]) and those in more downstream regions respond to more complex patterns of combined features [54, 55]. At first, neural coding studies primarily relied on retinotopy to infer visual content since the spatial organization of images is reflected in the stimulus-evoked responses in the primary visual cortex (V1) [56]. As such, visual content was mainly inferred from neural responses in early cortical areas, and the stimuli often consisted of low-resolution contrast patterns or digits [24, 25, 27, 29, 32]. Attempts to reconstruct more complex naturalistic images from activations in early regions were taken [28] but still fell short of capturing the full complexity of high-level neural activity required for reconstructing more intricate visual content. To successfully decode more high-level information from anterior regions, suitable feature representations were required that captured similar information about the stimulus as these responses, as attempted with more high-level hand-engineered features by [26] and [57] to reconstruct naturalistic images and scene backgrounds, respectively.

Next, the complexity gradient in visual processing, where increasingly complex features are represented across the ventral stream, was also identified in deep neural networks (DNNs): the alignment of DNN layers with neural activations revealed that early layers were mainly predictive of responses in upstream visual areas whereas deeper layers were more predictive of more downstream visual areas in humans [3, 4, 5, 6, 7, 8, 11] as well as in primates [1, 12]. At present, DNNs are commonly used to decode more high-level neural activity during visual perception, imagery and dreaming [58, 19, 33, 59, 34, 60, 35, 61, 36]. For reconstruction, the decoded feature representations of discriminative DNNs were used, for instance, by providing them directly as input to a decoder DNN (feature-to-image) [33] or by using a feature loss to iteratively optimize the pixel values in an input image [34] or decoder weights [62] so that the reconstruction features matched those of the stimulus. At this time, *unsupervised* learning paradigms, although more biologically plausible, seemed to appear less successful in modeling neural representations in the primate brain than their supervised counterparts [5].

Recent advancements have shifted attention towards the potential of *unsupervised generative* (rather than discriminative) models and their latent spaces, such as Variational Autoencoders (VAEs) [63, 35] and GANs [59, 64, 60, 61, 36]. In contrast to discriminative features, generative latents offer a distinctive advantage by aligning neural representations with generative processes the brain might perform during various cognitive functions (e.g., anticipation and mental imagery). Also, it is not possible to directly model the synthesis operation from discriminative features because they are primarily optimized to differentiate between classes rather than generate new visual content. However, the challenge posed by the scarcity of neural data, along with the substantial data requirements for properly training data-hungry DNNs with numerous parameters, has hindered effective GAN training from scratch (see [60], for an attempt using 6000 training examples). To address this issue, [59] trained an encoder model to generate synthetic neural activity to a much broader set of images which was then utilized to train a GAN. However, biases and inaccuracies in the synthetic neural data fail to capture the intricate details of authentic neural responses, leading to discrepancies between the reconstructions and stimuli. Rather than training our own models from limited data, we can also leverage pretrained GANs and their latent spaces as a proxy for brain activity. For this, access to the latents of the visual stimuli is required so that a linear model can be fit on these latents and the neural data, after which predicted latents from held-out brain activity can be fed to the GAN for image reconstruction. Yet, the inherent nonlinearities in the transformation from latent space to image space by the generator render it inherently unidirectional. Post-hoc approximate inversion has shown to work to some extent but entails information loss [64, 61] (note that VAEs do approximate inference by design). Instead, to have direct access to the ground-truth latents, [36] used synthesized stimuli by a pretrained progressively grown GAN, which was the state-of-the-art generative model for generating high-quality and high-resolution images at the time. The current work adopted and improved this experimental paradigm to study neural representations in the ventral visual stream during visual perception.

Finally, an earlier study already showed that disentangled latent units learned by a *β*-VAE better explained the coding of single neurons in the primate inferior temporal (IT) cortex at the end of the ventral stream during face perception [65]. This further underscores the potential of such generative models to unravel intricate neural representations and their interactions with complex visual stimuli.

## 2 Results

We used two datasets of visual stimuli. (i) Face images synthesized by StyleGAN3 (pretrained on the Flickr Faces High-Quality (FFHQ) dataset) consisting of 4000 and 100 training and test set images, respectively. (ii) High-variety natural images synthesized by StyleGAN-XL (pretrained on ImageNet), consisting of 4000 and 200 training and test set images, respectively.

### 2.1 Neural encoding

We studied how well neural responses were predicted from latents of recent generative models. Specifically, we focused on three types of latents: *z*-latents of StyleGAN3/StyleGAN-XL (512-/128-dim.), feature-disentangled *w*-latents of StyleGAN3/StyleGAN-XL (512-/512-dim.) and language-regularized CLIP-latents (768-dim.). In the case of natural images, we used the embedding that integrated both *z*-latent and class information, which serves as the input for the first layer of the mapping MLP. For each individual unit within a multi-unit microelectrode (960 individual units in total), we fit three distinct kernel ridge regression models on the aforementioned *z*-, *w*- and CLIP latents, of which the optimal regularization parameter *λ* was determined per visual area using 5-fold cross-validation.

For reference, we also fit three distinct encoding models on feature representations extracted from the discriminative VGG16 network, which was pretrained for either face or object recognition. Concretely, we used early (1; layer 2/16, after max pooling), middle (2; layer 7/16, after max pooling) and deep (5; layer 13/16, after max pooling) activations for this purpose. Note that the numbering system ’1, 3, 5’ refers to the max pooling operations in VGG16, which has a total of five max pooling layers. This numbering is used for the remainder of this manuscript. The performance of the six encoding models, per image dataset, was quantified by Pearson product-moment correlation coefficients (Figure 5). As discussed in ’earlier work’, the complexity gradient observed across the ventral stream in the brain is also reflected in the multi-layered architecture of discriminative DNNs [3, 4, 5, 6, 7, 8, 11, 1, 12]. As such, the representations extracted from early layers are more predictive of responses in early visual areas, while the deeper representations are more predictive of responses in more downstream areas. We reproduced this complexity gradient by assigning the discriminative representation with the highest encoding performance to each microelectrode unit on the brain (Figure 8, first graph).

**Figure 5:**
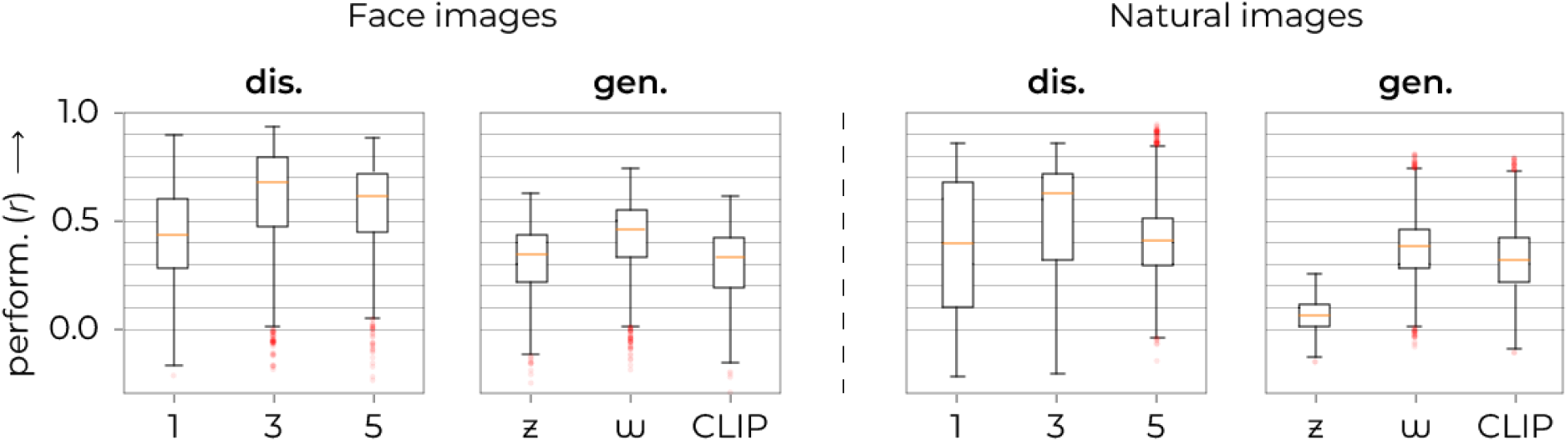
Encoding performance. The effectiveness of each encoding model is assessed using the Pearson correlation coefficients between predicted and recorded neural responses. For each dataset, the first and second graphs denote discriminative and generative representations, respectively. The correlation distribution across each encoding model shows a robust level of accuracy.

**Figure 6:**
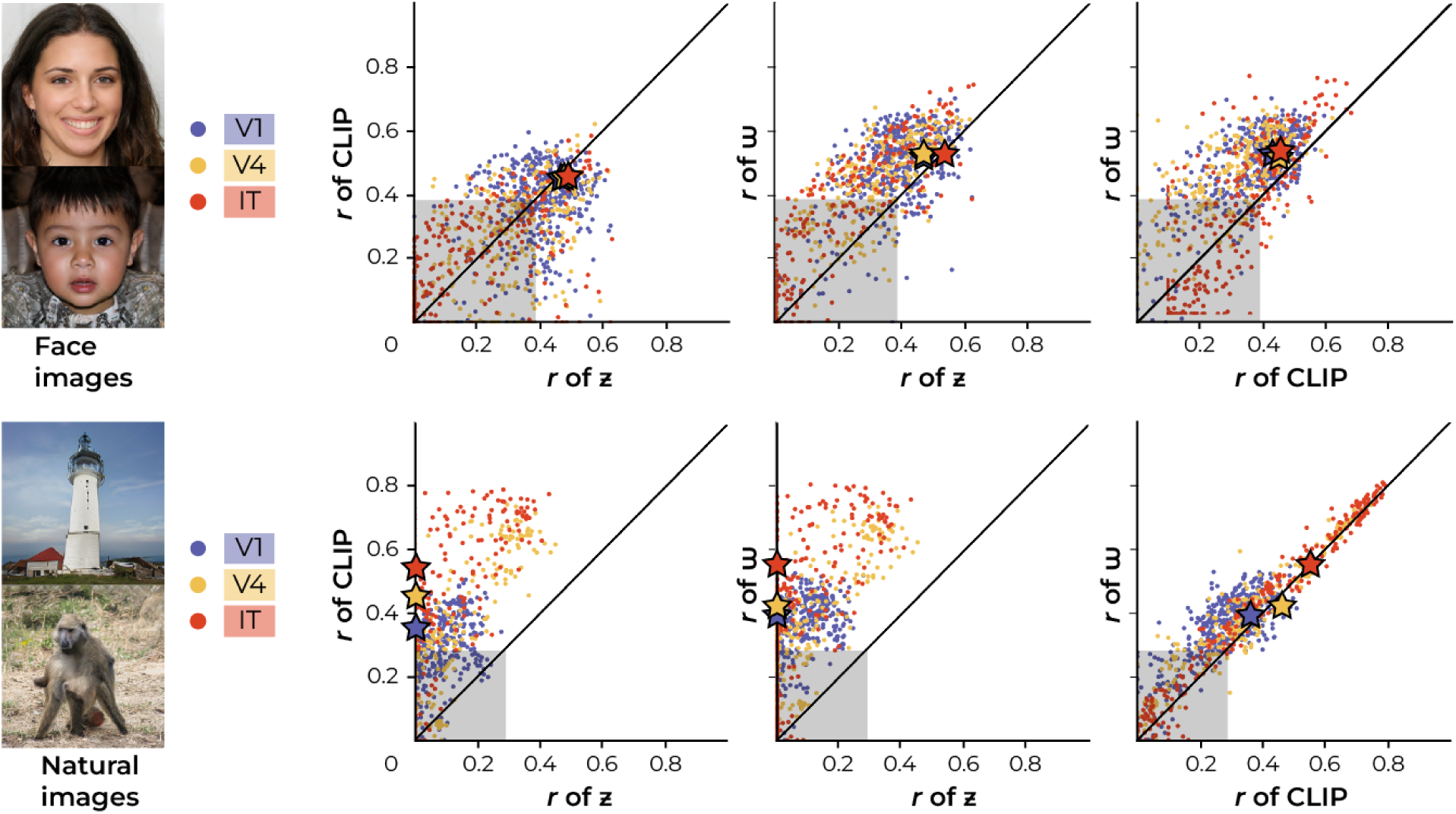
Generative-based encoding performance. For each individual microelectrode unit, we fit three encoding models based on three distinct feature representations: *z*-, *w−* and CLIP-latent representations. As such, we fit 3*×* 960 independent encoders, resulting in 3*×* 960 predicted neural responses because there were seven, four and four microelectrode arrays (64 units each) for V1, V4 and IT, respectively (i.e., 7 *×* 64 = 448 in V1, 4 *×* 64 = 256 in V4 and 4 *×* 64 = 256 in IT). The scatterplots display the prediction-target correlation (*r*) of one encoding model on the X-axis and another encoding model on the Y-axis to investigate the relationship between the two. Each dot represents the performance of one modeled microelectrode unit in terms of both encoding models (so, 960 dots per plot). Negative correlation values were set to zero. The diagonal represents equal performance between both models. The critical *r*-value at Bonferonni-corrected *α* = 5.21e*−*5 is at *r* = 0.3895 and *r* = 0.2807 for faces (*df* = 100) and natural images (*df* = 200), respectively, and is denoted by the shaded area. It is clear to see that *w*-latents outperform both *z*- and CLIP-latents because most dots lie in the direction of the *w*-axis (above the diagonal). The stars indicate the mean correlation coefficient per region of interest based on the data points outside the shaded area.

Notably, among the generative-based encoders, the *w*-latent-based encoder statistically outperformed those of *z*- and CLIP-latents in predicting the neural activity (Figure 5, 6). For face images, the *w*-latent-based encoder demonstrated significant superiority over the *z*-based encoder (2-Sample T-Test; *t*(1918) = *−*13.8067, *p* = 2.07e*−*41) as well as the CLIP-based encoder (2-Sample T-Test; *t*(1918) = 16.0527, *p* = 1.65e*−*54). Additionally, the CLIP-based encoding also outperformed the *z*-based encoding (2-Sample T-Test; *t*(1918) = 2.1068, *p* = 0.0353) although the difference was not as pronounced. Similarly, for natural images, the *w*-latent-based encoder significantly outperformed the *z*-based encoder (2-Sample T-Test; *t*(1918) = *−*44.4495, *p* = 3.13e*−*297) and the CLIP-based encoder (2-Sample T-Test; *t*(1918) = 6.2957, *p* = 3.78e*−*10). And CLIP-based encoding also outperformed *z*-based encoding (2-Sample T-Test; *t*(1918) = *−*35.3777, *p* = 1.79e*−*211). Figure 7 directly compares the raw *w*-based encoding performance across visual areas and shows that the *w*-latents of natural images mainly captured visual information relevant to high-level neural activity, as indicated by the increasing variance explained from V1 to IT. This pattern is however not observed for face images.

**Figure 7:**
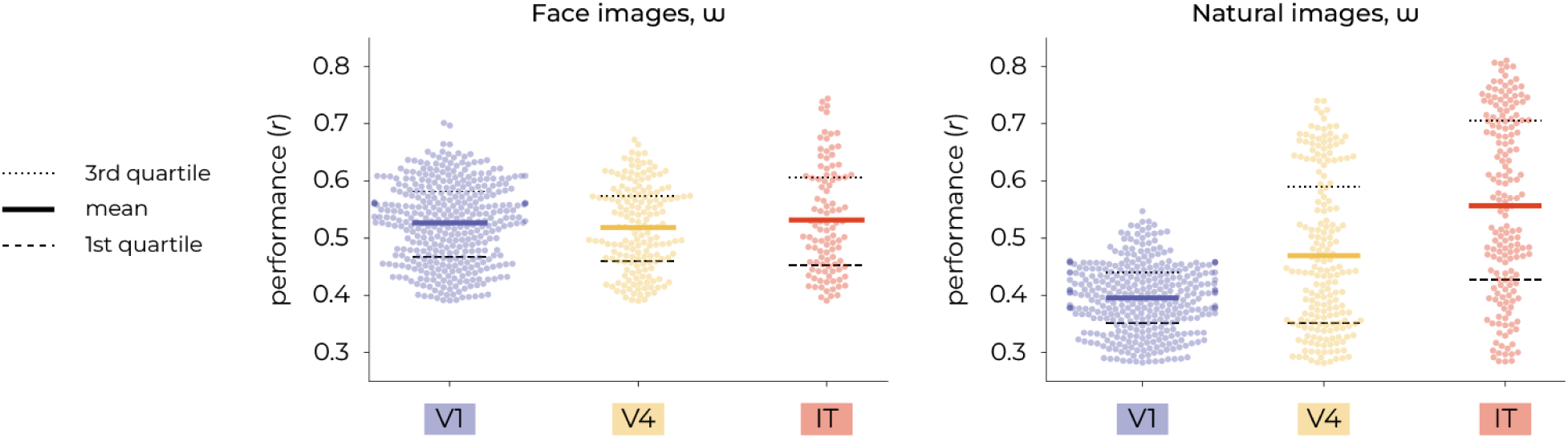
*w*-based encoding performance across visual areas. The left panel presents the distribution of correlation coefficients for face images using a swarm plot, with mean values indicated for V1 (0.53), V4 (0.52), and IT (0.53). The right panel displays the distribution for natural images, with mean values for V1 (0.40), V4 (0.47), and IT (0.56).

Subsequently, we also explored the positioning of generative *w*-latents along the complexity gradient (originally defined by discriminative features) which refers to the progression from simpler, lower-level visual processing in early visual areas to more complex, higher-level processing in areas like IT. To this end, we replaced each level of discriminative feature representation with the *w*-latent representation to see where along this gradient the *w*-latents have the most predictive power (Figure 8). The results of this comparative analysis revealed that *w*-latents of both image types are predominantly assigned to the higher end of the complexity spectrum. This indicates that *w*-latents capture visual features that are particularly relevant to high-level neural activity. This positioning should not be interpreted as a competition between discriminative and generative latents; rather, it highlights their complementary nature as high-level representations in the overall hierarchy of neural encoding. It is possible that while *w*-latents explain more variance in higher visual areas for both face and natural images (as seen in Figure 8), the increase in variance explained from V1 to IT (the gradient) is more pronounced for natural images than for face images (as suggested by Figure 7) and therefore not as apparent when looking at *w*-latents in isolation.

**Figure 8:**
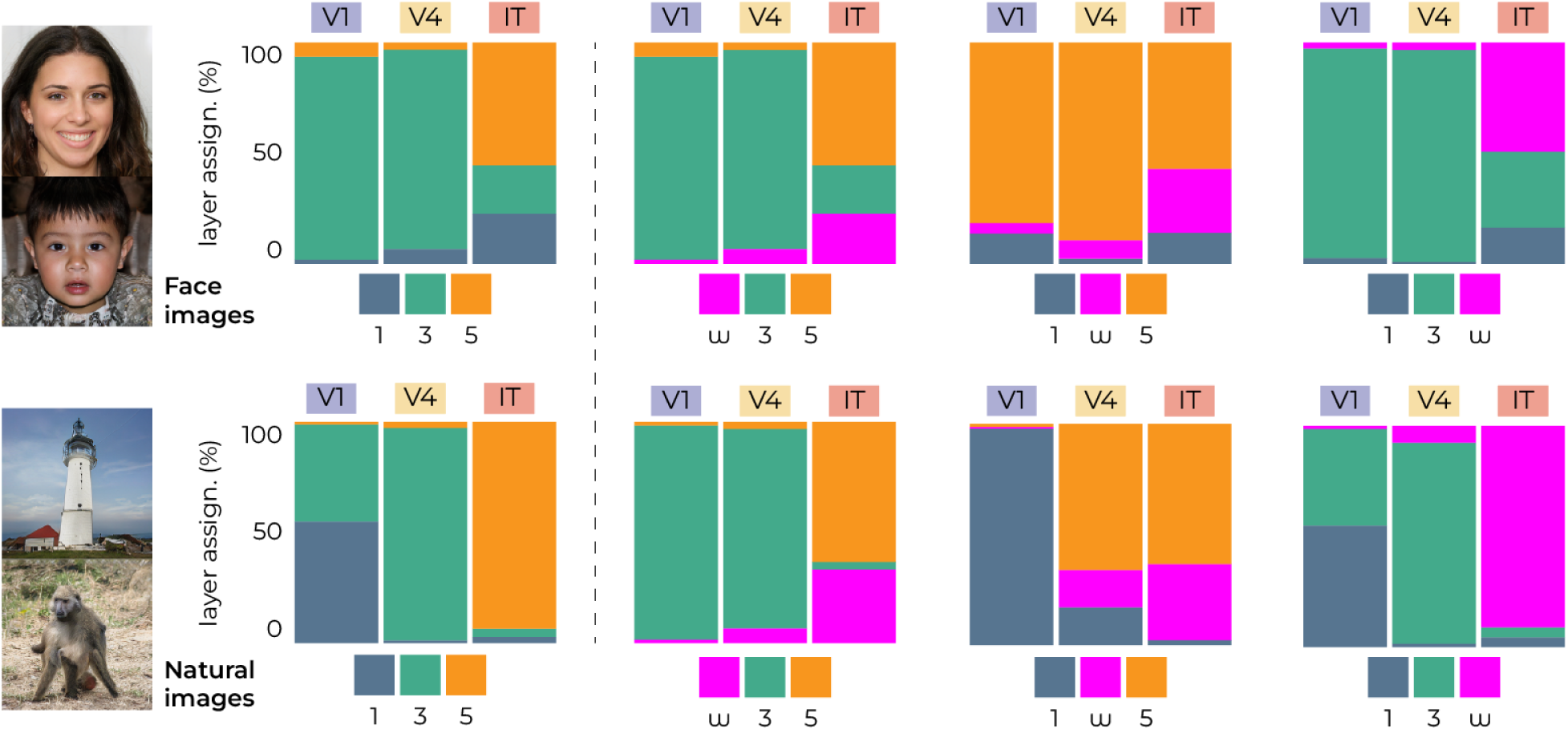
*w*-latents explain high-level brain activity. Three encoding models were fit on early (1; layer 2/16), middle (3; layer 7/16) and deep (5; layer 13/16) feature representations of VGG16 pretrained for face/object recognition. The representation that led to the highest encoding performance was assigned to each microelectrode unit, resulting in the complexity gradient where more low-level and high-level representations are assigned to earlier and more downstream brain areas, respectively (see most-left graph for reference). In each of the three plots, one VGG16 representation was replaced by the *w*-latent representation to see where it falls on the complexity gradient. The results illustrate that *w*-latents predominantly accounted for neural responses in downstream IT.

It is also worth noting that encoders based on discriminative models appear to generally outperform those based on generative models. The performance proximity of *w*-latents to some discriminative-based predictions suggests that feature disentanglement in generative models may enhance their predictive capabilities, but they inherently differ from discriminative models in their primary function and approach to data. However, given that the *w*-based and discriminative-based encoders are compared in Figure 8, we statistically analyzed their comparisons for a more informed understanding of the relative strengths of each encoding approach. For faces, the comparison between *w*-based and VGGFace-1-based encoders shows no significant difference (2-Sample T-Test; *t*(1918) = *−*0.2859, *p* = 0.78) but we found highly significant differences when comparing *w*-based encoders with VGGFace-3 (2-Sample T-Test; *t*(1918) = 16.5817, *p* = 8.21*e −* 58) and with VGGFace-5 (2-Sample T-Test; *t*(1918) = 15.0820, *p* = 1.17*e −* 48). For natural images, the disparity between *w*- and VGGFace-1-based encoders also shows no significance (2-Sample T-Test; *t*(1918) = 0.7771, *p* = 0.4372). In contrast, we observed a highly significant difference between between *w*-based and VGGFace-3-based encoders (2-Sample T-Test; *t*(1918) = 12.7855, *p* = 5.56*e −* 36) and a significant difference between *w*-based and VGGFace-5-based encoders (2-Sample T-Test; *t*(1918) = 3.7425, *p* = 0.0002). Despite their statistical similarity to the encoders based on early activations of VGG16-1, *w*-based encoders are mainly predictive of high-level brain activity in IT.

### 2.2 Neural decoding

The ‘analysis‘ component of neural decoding was modeled by multiple linear regression from neural responses to the feature-disentangled *w*-latents, which were subsequently fed to the generator for ‘synthesis‘. This resulted in remarkably accurate reconstructions that closely resembled the stimuli in their specific characteristics; Figure 9, 10. Perceptually, we can notice high similarity between stimuli and their reconstructions in terms of their specific attributes (e.g., gender, age, pose, haircut, lighting, hair color, skin tone, smile and eyeglasses for faces; shapes, colors, textures, object locations, (in-)animacy for natural images). We repeated the experiment with another macaque that had silicon-based electrodes in V1, V2, V3 and V4 (Appendix S1).

**Figure 9:**
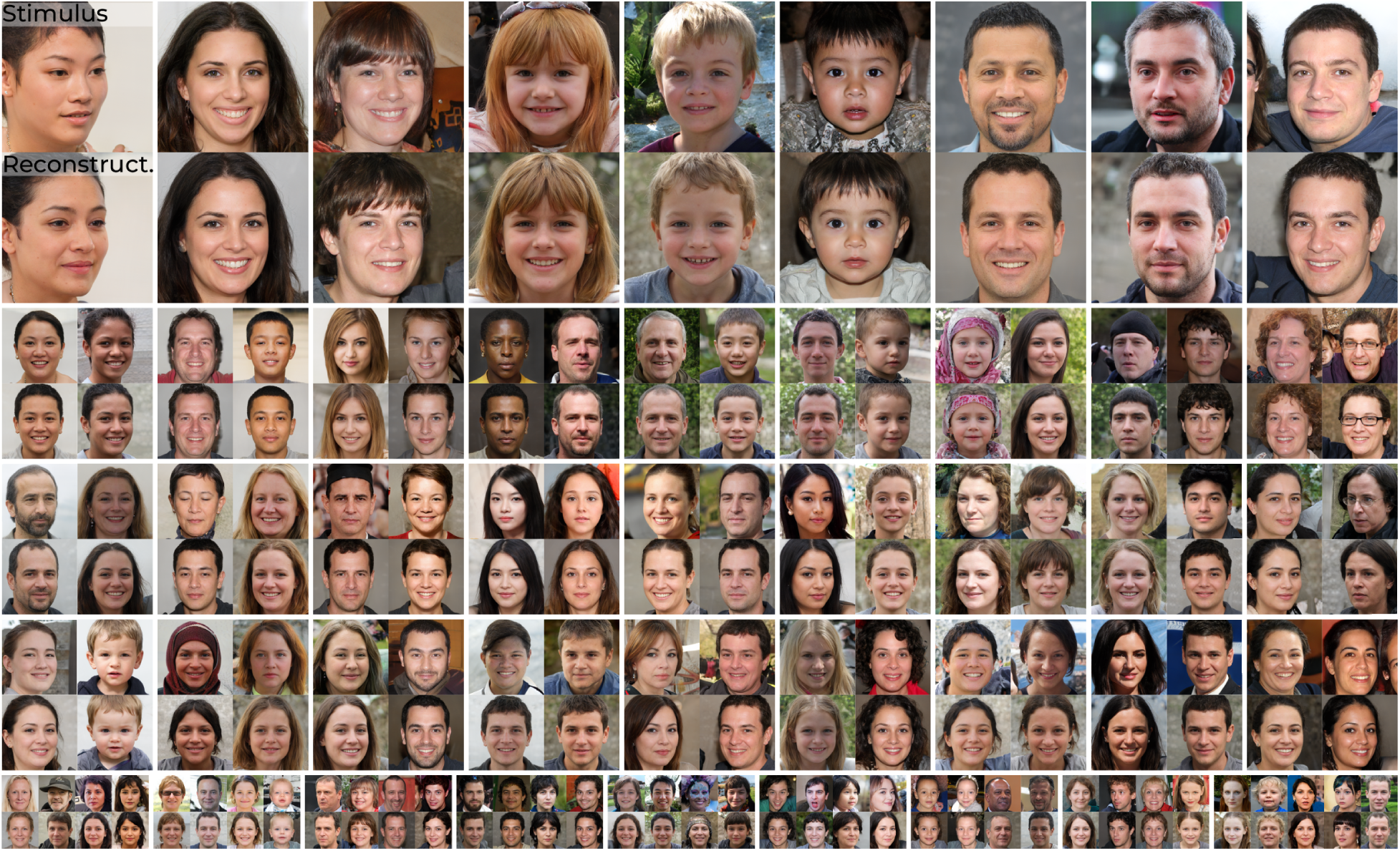
Qualitative reconstruction results: The 100 test set stimuli (top row) and their reconstructions from brain activity in V1, V4 and IT (bottom row) via *w*-latents.

**Figure 10:**
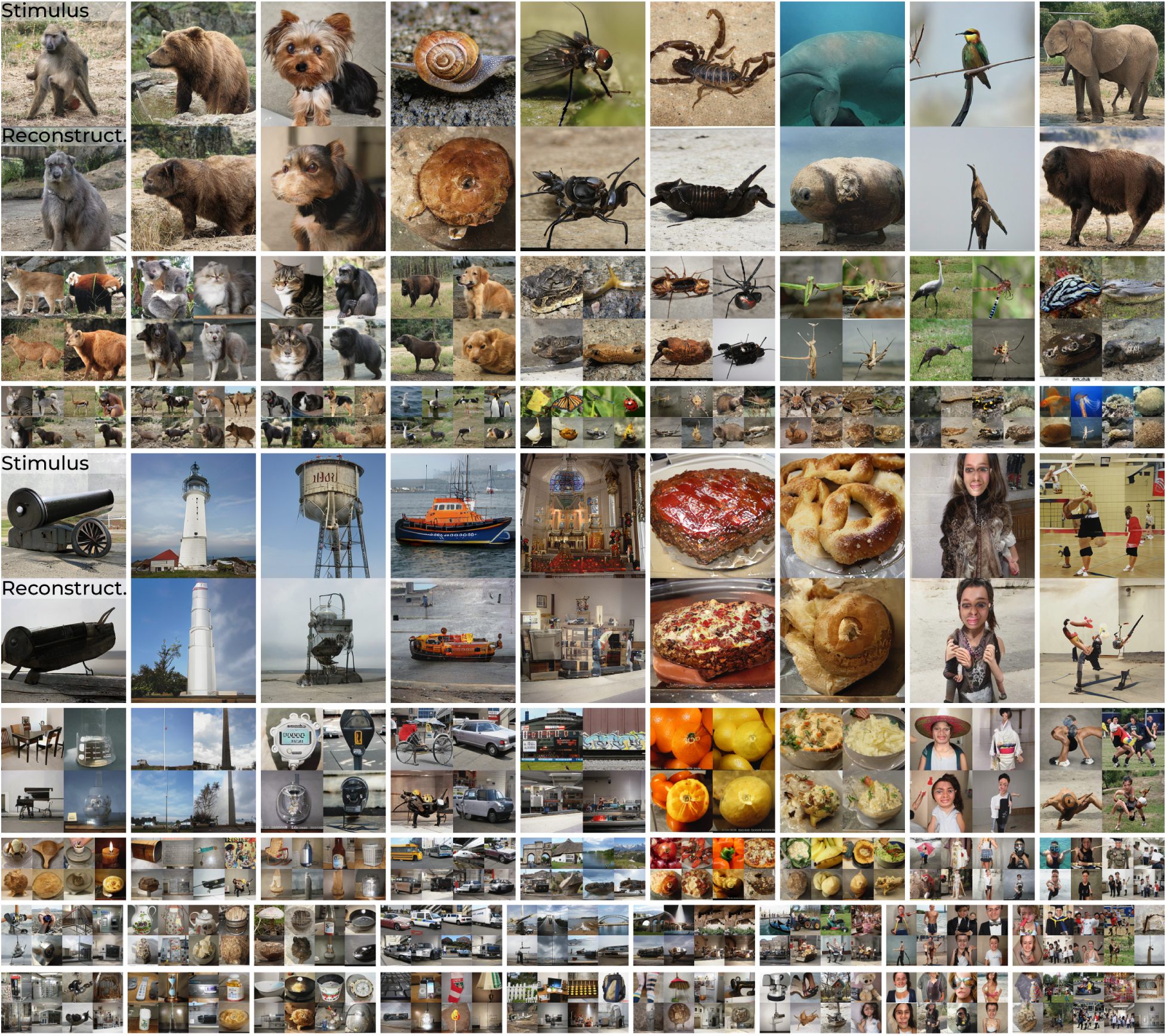
Qualitative reconstruction results: The 200 test set stimuli (top row) and their reconstructions from brain activity in V1, V4 and IT. (bottom row) via *w*-latents.

The supplementary materials contain decoding results from *z*-latents (Appendix S2) and another reconstruction approach based on [28] (Appendix S3). The former not only demonstrates superior performance using *w*-latents over *z*-latents in *conditional* image generation but also that this disentanglement enables *unconditional* image generation using GANs. Furthermore, a leave-one-class-out analysis confirmed that our approach extends beyond mere classification (Appendix S5).

The quantitative metrics in Table 1 show the similarity between stimuli and their reconstructions from brain activity in terms of six metrics that evaluated reconstruction quality at different levels of abstraction (see Appendix S7 for the visual guide). Specifically, a stimulus and its reconstruction were both fed to VGG16 (pretrained on face- and object recognition for faces and natural images, respectively) and we extracted five intermediate activations (the five MaxPool layers) thereto. The early layers capture more low-level features (e.g., edges and orientations) whereas deeper layers capture increasingly higher-level features (e.g., textures to object parts to entire objects). We then compared the cosine similarity between these extracted representations of stimulus and reconstruction. Next, to study the decoder that resulted in these accurate reconstructions, the contribution of each visual area was determined by the occlusion of the microelectrode recordings in the other two brain areas (rather than fitting three independent decoders on subsets of brain activity). It is reasonable to say that, of the three cortical areas, the area that resulted in the highest similarity contains the most information about that representation. For faces, decoding performance was for the largest part determined by responses from IT - which is the most downstream site we recorded from. For natural images, we found that the lower-level representations (VGG16 layers 1-2) were most similar when decoded from V1 and the higher-level representations (VGG16 layers 3-5) and latent space were most similar when decoded from area IT. We validated our quantitative results with a permutation test as follows: per iteration, we sampled a hundred/two-hundred random latents from the same distribution as our original test set and generated their corresponding images. We assessed whether these random latents and images were closer to the ground-truth latent and images than our predictions from brain activity, and found that our predictions from brain activity were always closer to the original stimuli than the random samples for all metrics, yielding statistical significance (*p <* 0.001) (in Appendix S6, the results of random permutation analyses can be found).

**Table 1:** Quantitative results. The upper and lower block display model performance (*mean ± std.error*) when reconstructing face images and natural images, respectively, in terms of six metrics of perceptual cosine similarity using the five MaxPool layer outputs of VGG16 for face recognition (face images) / object recognition (natural images) and latent cosine similarity between *w*-latents of stimuli and their reconstructions. The rows display decoding performance when using the recordings from all recording sites (i.e., V1, V4 and IT together) or the recordings within a specific brain area.

|  |  | VGG16-1 sim. | VGG16-2 sim. | VGG16-3 sim. | VGG16-4 sim. | VGG16-5 sim. | Lat. sim. |
| --- | --- | --- | --- | --- | --- | --- | --- |
| Face images | all | **0.7871 ± 0.0102** | **0.7681 ± 0.0075** | **0.5874 ± 0.0075** | **0.6170 ± 0.0085** | **0.5940 ± 0.0104** | **0.5548 ± 0.0045** |
| | V1 | 0.6382 ± 0.0079 | 0.6758 ± 0.0064 | 0.4891 ± 0.0064 | 0.5041 ± 0.0083 | 0.4442 ± 0.0092 | 0.5022 ± 0.0047 |
| | V4 | 0.6303 ± 0.0101 | 0.6729 ± 0.0068 | 0.4890 ± 0.0068 | 0.5006 ± 0.0085 | 0.4191 ± 0.0091 | 0.5026 ± 0.0040 |
| | IT | 0.7123 ± 0.0110 | 0.7133 ± 0.0073 | 0.5093 ± 0.0073 | 0.5253 ± 0.0087 | 0.4434 ± 0.0096 | 0.5176 ± 0.0039 |
| Natural images | all | **0.4083 ± 0.0036** | **0.3322 ± 0.0036** | **0.2555 ± 0.0025** | **0.2192 ± 0.0043** | **0.2497 ± 0.0066** | **0.8032 ± 0.0032** |
| | V1 | 0.3929 ± 0.0031 | 0.3147 ± 0.0031 | 0.2223 ± 0.0019 | 0.1511 ± 0.0023 | 0.1367 ± 0.0037 | 0.7336 ± 0.0036 |
| | V4 | 0.3790 ± 0.0029 | 0.3132 ± 0.0029 | 0.2270 ± 0.0019 | 0.1641 ± 0.0027 | 0.1617 ± 0.0045 | 0.7614 ± 0.0034 |
| | IT | 0.3798 ± 0.0026 | 0.3127 ± 0.0026 | 0.2302 ± 0.0020 | 0.1790 ± 0.0039 | 0.1692 ± 0.0057 | 0.7653 ± 0.0039 |

#### 2.2.1 Time-based neural decoding

Time-based neural decoding showed the gradual extraction of stimulus-related information over the trial of 300 ms, with stimulus presentation occurring at 100 ms, by sliding a 100 ms time window across the entire time course using a stride of 25 ms, resulting in nine averaged points of neural activity across time (Figure 11A). We fit separate decoders for individual time points but decoding via the original decoder, which was fit on brain activity within the predefined time windows, yielded similar results. Initially, the reconstructions exhibited an average appearance, but then gradually acquired their distinct visual features upon stimulus onset (Figure 11B, D). Noteworthy, the reconstructions prior to stimulus onset exhibit an average-looking appearance because we averaged multiple repetitions in the test set, where each repetition was preceded by a different stimulus due to the randomized order of stimulus presentation. Although canceled out following our approach, it remains highly plausible that the information about the preceding stimulus is still preserved in the brain. Moreover, the area-based reconstructions and performance graphs revealed that V1 generally displayed stimulus-like visual features earlier in time whereas IT consistently outperformed the other two in the final reconstruction of stimulus information (Figure 11C, E). Albeit trivial, the finding that reconstruction from all rather than isolated areas yields the highest performance confirms that visual perception involves a distributed process across multiple areas that each hold distinct information about the stimulus.

**Figure 11:**
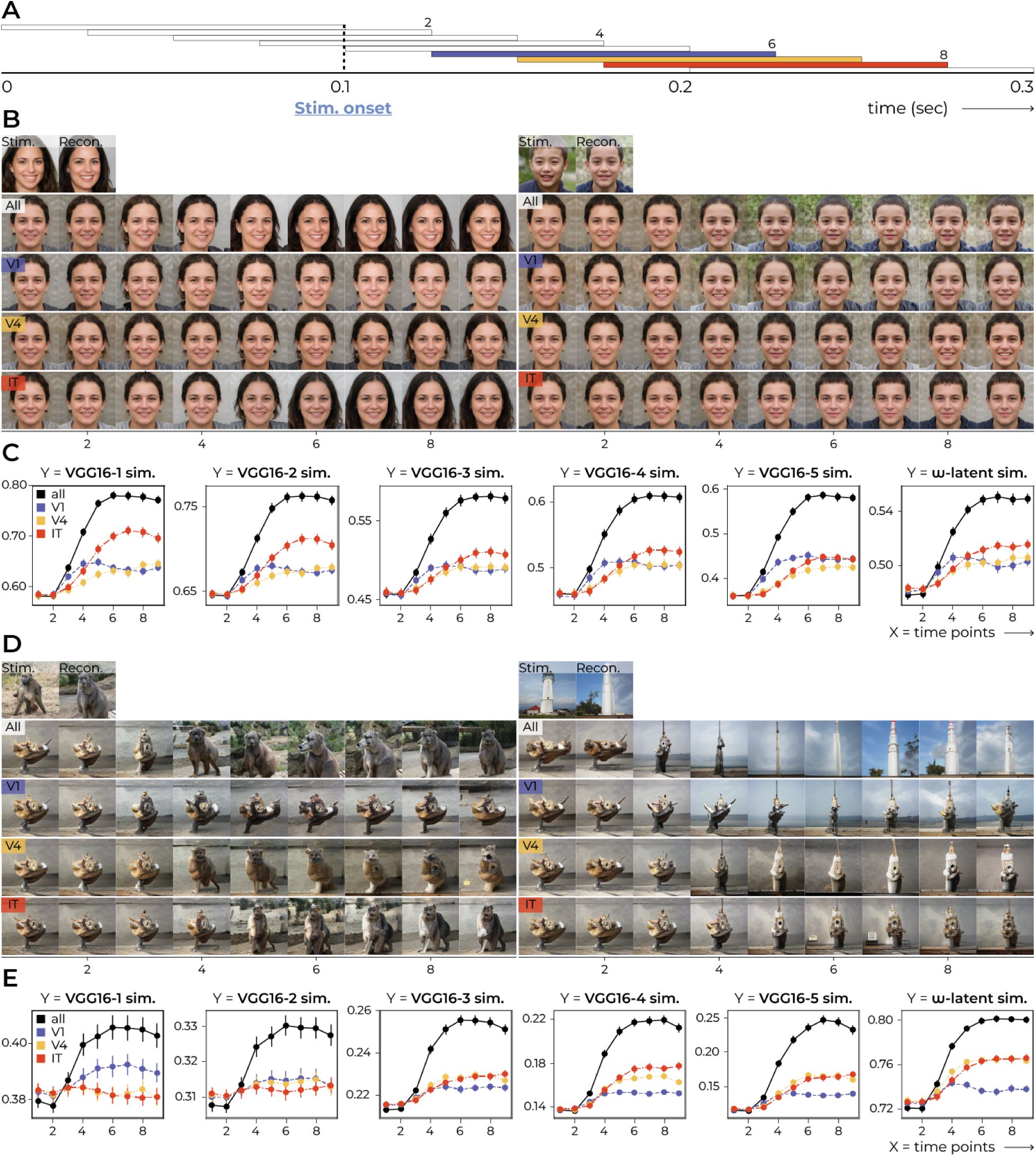
Time-based decoding. **A** For each trial, responses were recorded for 300 ms with stimulus onset at 100 ms. Rather than taking the average response within the *original* time windows (see the three color-coded windows for V1, V4 and IT), we slid a 100 ms window with a stride of 25 ms over the entire time course, resulting in nine average responses across time. **B, D** Two stimulus-reconstruction examples evolve over time for faces and natural images, respectively. **C, E** Decoding performance over time for faces and natural images, respectively. The error bars denote the standard error of the cosine similarities between features of stimuli and reconstructions. It can be noted how V1 performance climbs up slightly earlier in time than the other two visual areas. For faces, IT outperforms V1 and V4 in most instances. For natural images, V1 outperforms V4 and IT for low-level feature similarity, after which V4 and IT climb up together and outperform V1 for the more high-level feature similarity metrics.

#### 2.2.2 Linear operations

The application of linear operations to GAN-latents directly translates to meaningful perceptual changes in the generated images because visual data that look perceptually similar in terms of certain features are also closely positioned in latent space. As such, pathways through the well-structured latent landscape can be explored by interpolating two distinct latents, resulting in an ordered set of images whose semantics vary smoothly with latent codes [66] (Figure 12A1, B) and simple arithmetic operations [67] (Figure 12A2). Since such operations can be performed to traverse the latent space (Figure 12, row 1), without needing to understand the intricate details of the underlying generator network, the latent-response correspondence also opens the door to interpreting neural activity in terms of such operations within the latent space (Figure 12, row 2). To illustrate, consider having a neural response to a neutral face and another neural response to a smiling face, interpolating their respective decoded latents yields a sequence of latents, and consequently, a series of images transitioning from neutral to smiling expressions. Note that both the operations applied to neural activity as well as the decoder are linear, resulting in “linearity stacking”. This means that we can also apply the linear operations directly to the neural responses themselves, decode them into latents, and feed them to the GAN for reconstruction. This would yield the same images as those in row 2 of Figure 12.

**Figure 12:**
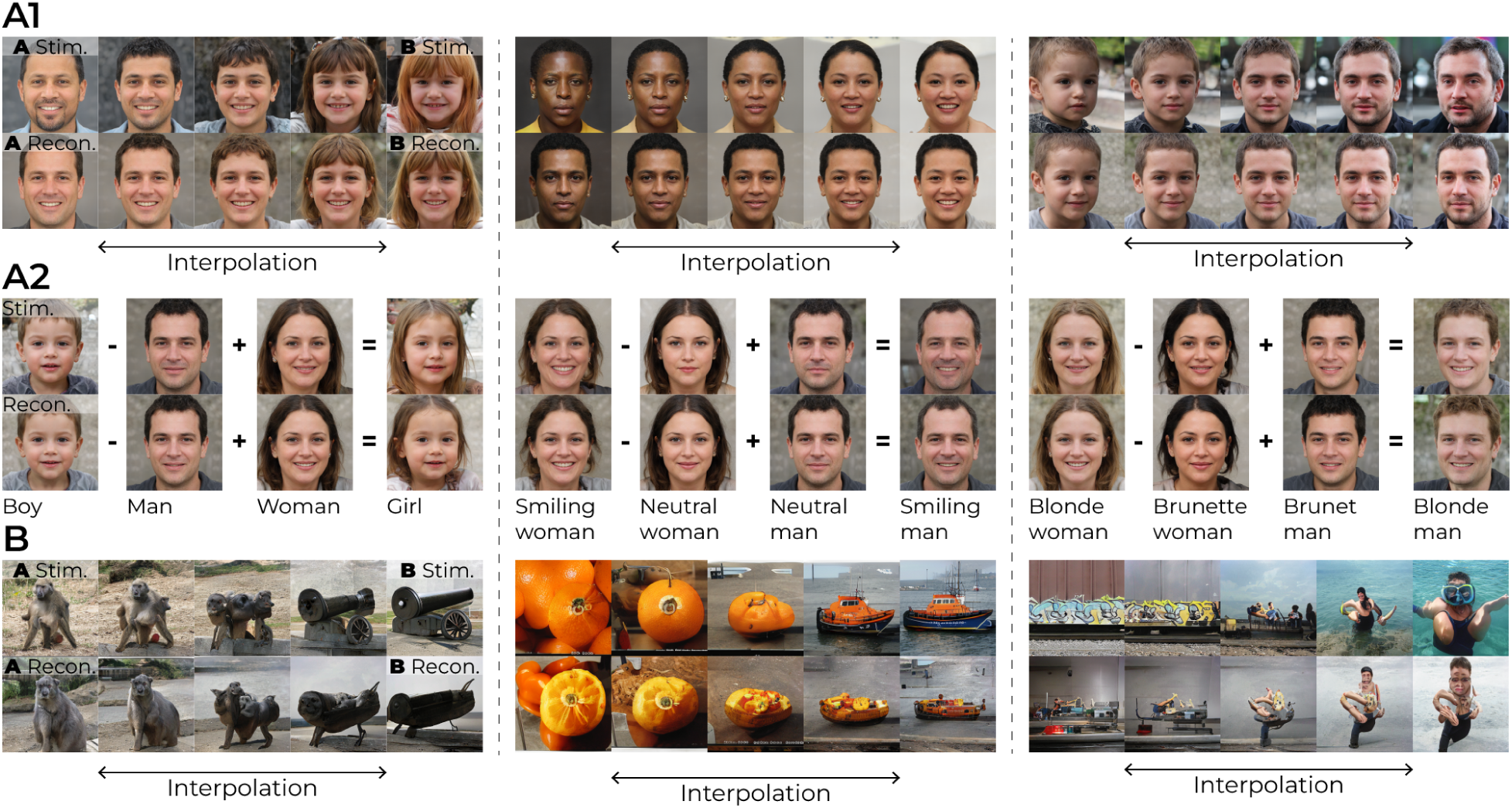
Linear operations to latent codes. *(row 1)* shows linear operations to two ground-truth *w*-latents, and *(row 2)* to two predicted *w*-latents from brain activity. The linearly-manipulated latents were then fed to the generator for image generation. (A1, A2) face images, also contains vector arithmetic. (B) As for (A1, A2) but for natural images.

The linear relationship between neural activity patterns and latent codes, coupled with the feature-disentangled nature of the GAN’s latent space, enables synthesis (and analysis) of specific aspects of the visual experience captured by the neural responses - which is the key idea of our decoding approach.

## 3 Discussion

In this study, we characterized neural representations of visual perception using high-level latent representations of generative models. Our encoding analysis showed feature-disentangled *w*-latents conditioned on StyleGAN3/StyleGAN-XL to outperform the other latent candidates in explaining neural responses. Subsequently, we used the *w*-latents for neural decoding of the recorded brain activity, which resulted in reconstructions that strongly resembled the original stimuli in their specific characteristics. Given the virtually infinite number of possible candidate representations to encode the same image, finding a representation that accurately reflects the information in brain activity is not a trivial task. In our approach, the decoded *w*-latents resulted in image reconstructions that closely matched the stimuli in their semantic as well as structural features. Overall, this work highlights the importance of feature disentanglement in explaining high-level neural responses and demonstrates the potential of aligning such unsupervised generative models with biological processes. These findings have implications for the advancements of computational models and the development of clinical applications for people with disabilities. For instance, neuroprosthetics to restore vision in blind patients as well as brain-computer interfaces (BCIs) to enable nonmuscular communication with individuals who are locked in.

### 3.1 Uncovering principles of neural coding

The primary goal of our study was to uncover principles that govern neural coding of the visual world and gain a more interpretable understanding of high-level neural representations underlying visual perception using deep generative modeling. As such, similarities between *w*-latents and the brain could provide further insights into what drives the organization of visual processing in the brain. First, GANs are trained in an unsupervised setting; they learn directly from raw visual data without explicit labels or annotations. Not only does this make GANs more biologically plausible than their supervised counterparts since it resembles more closely how the brain learns from its environment but they may also lead to more flexible and generalizable representations that are better able to capture the underlying structure and patterns in the observed data. Note that our finding that discriminative-based encoders (supervised) outperform *w*-based encoders (unsupervised) in neural encoding does not directly challenge this notion since these models were optimized for different objectives (i.e., image recognition and image generation, respectively). Second, StyleGAN was designed to disentangle different visual semantics into separate *w*-latent features. The superior performance of *w*-latents relative to other generative latents highlights the role of feature disentanglement in explaining the high-level neural representations, and the ability to disentangle the object manifold [68]. Keep in mind that StyleGAN itself has never been optimized on neural data which implies a general principle of shared encoding of real-world phenomena. Finally, there is a conceptual analogy between the adversarial training of GANs and the predictive coding theory of perception where the brain uses top-down predictions, based on prior knowledge and experience, to guide bottom-up sensory processing and adjusts its internal models based on the mismatch between expectations and actual observations. In GANs, the discriminator and generator engage in a similar process with the discriminator evaluating the “real” sensory input and the “predicted/imagined” instances by the generator. Based on the mismatch, as determined by the discriminator, the internal model of the generator is refined such that its outputs match the real-world data closer; the generator harnesses the knowledge of the discriminator to learn how to represent the world in its latent space. And like generative models interpolate between latent vectors to create intermediate outputs, the brain might engage in similar processes that interpolate between neural representations to accommodate variations in mental simulation. So, while the exact mechanisms used by the brain and GANs differ significantly, their conceptual similarities could provide insights into the nature of perception and the potential of machine learning to capture some of the same principles underlying this ability.

### 3.2 Limitations and future directions

It is essential to clarify that the brain’s overall functioning is far more complex than a linear system; our approach merely exploits the linearity within a particular representation space. Further investigation of the correspondence between latents and responses is needed by, for instance, obtaining neural responses to more diverse stimuli and stimulus manipulations to identify which visual properties could be effectively translated to the latent space and where this approach falls short. We did observe that, within the limitations of StyleGAN-XL’s design tied to the natural image distribution, the generator exhibits the capability to synthesize abstract stimuli (see Appendix S8), which offers a promising perspective for future investigations in this direction. Further, this study solely used synthesized stimuli with known latent representations generated by StyleGAN. While this allowed for a controlled and systematic examination of neural representations of visual information, future studies should also include real photographs to see how this method generalizes. This requires accurate inversion methods of the generator’s ‘synthesis‘ operation, yet this endeavor is intricate due to the inherent information loss associated with post-hoc inference. That said, the current study still performed valid neural encoding and reconstruction from brain activity despite the nature of the presented images themselves. Another limitation is the small sample size of one subject (note that we did include face reconstructions from a second subject with different cortical implants in Appendix S1). Although small sample sizes are common in studies using invasive recordings - larger sample sizes are needed to further confirm the robustness of our findings. Finally, it is worth noting that the use of deep neural networks to model brain activity is still a developing field and the models used in this study are not flawless representations of the underlying neural processes.

## 4 Materials and Methods

### 4.1 Stimuli

StyleGAN [41, 69] was developed to optimize control over the semantics in the synthesized images in single-category datasets (e.g., only-faces, -bedrooms, -cars or -cats) [42]. This generative model maps *z*-latents via an MLP to an intermediate *w*-latent space in favor of feature disentanglement. That is, the original *z*-latent space is restricted to follow the data distribution that it is trained on (e.g., old-looking faces wear eyeglasses more often than young-looking faces) and such biases are entangled in the *z*-latents. The less entangled *w*-latent space overcomes this such that unfamiliar latent elements can be mapped to their respective visual features.

**Dataset i: Face images.** We synthesized photorealistic face images of 1024 *×* 1024 px resolution from (512-dim.) *z*-latent vectors with the generator network of StyleGAN3 (Figure 3) which is pretrained on the high-quality Flickr-Faces-HQ (FFHQ) dataset [41]. The *z*-latents were randomly sampled from the standard Gaussian. We specified a truncation of 0.7 so that the sampled values are ensured to fall within this range to benefit image quality. During synthesis, learned affine transformations integrate *w*-latents into the generator network with adaptive instance normalization (like *style transfer* [70]). Finally, we synthesized a training set of 4000 face images that were each presented once to cover a large stimulus space to fit a general model. The test set consisted of 100 synthesized faces.

**Dataset ii: Natural images.** Recently, StyleGAN-XL (three times larger in depth and parameter count than a standard StyleGAN3) was developed to scale up to larger and less-structured datasets using a new training strategy [71]. Concretely, the new training strategy combined (i) *the progressive growing paradigm* where architecture size is gradually increased by adding new layers, (ii) *the projected GAN paradigm* where both synthesized and real samples are mapped to four fixed feature spaces before being fed to four corresponding and independent discriminator networks, and (iii) *classifier guidance* where the cross-entropy loss of a pretrained classifier is added as a term to the generator loss. As such, StyleGAN-XL has been successfully trained on ImageNet [72] to generate high-resolution images of a thousand different categories, resulting in a complex and diverse stimulus dataset. We synthesized images from the 200 classes from Tiny ImageNet (a subset rather than all thousand classes from ImageNet)±[73] so that each class was represented by twenty training set stimuli and one test set stimulus (Appendix S9 lists the labels). First, a 64-dimensional vector was sampled from a standard Gaussian and concatenated with the 64-dimensional embedded representation of the class category, resulting in 128-dimensional *z*-latents that were utilized to synthesize 512 *×* 512 px resolution RGB images. For the training set, *z*-latents were randomly sampled and mapped to *w*-latents that were truncated at 0.7 to support image quality as well as diversity. The average *w*-latent of each category was utilized for the test set due to the high quality and because variation was not required as we only used one image per category (in Appendix S4, we qualitatively confirmed that our findings were not attributed to the use of the average w-latent). The *z*-latents of the test set were obtained by activation maximization of an input vector by minimizing its distance to the target *w*-latent. In total, the training and test set consisted of 4000 (each presented once) and 200 stimuli (averaged over 20 repetitions), respectively.

### 4.2 Features

As the in-between feature candidates, we used the (generative) *z*-latents of StyleGAN3/StyleGAN-XL (512-/ 128-dim.), *w*-latents of StyleGAN3/StyleGAN-XL (512-/512-dim.), and CLIP-latents (768-dim.). In the case of natural images, we used the embedding that integrated both *z*-latent and class information, which serves as the input for the first layer of the mapping MLP. We also used the five (discriminative) layer activations of VGG16 for face recognition [74] and object recognition [75]. Specifically, we utilized the outputs from layers 2/16, 4/16, 7/16, 10/16, 13/16, referred to as layers 1-5, following max pooling. Because the features from layer 1 and 2 were very large (*∼* 10^6^), we performed downsampling, as done in [11]. That is, for each channel in the activation, the feature map was spatially smoothed with a Gaussian filter and subsampled with a factor 2. The kernel size was set to be equal to the downsampling factor.

### 4.3 Responses

We recorded multi-unit activity (MUA) [47] with 15 chronically implanted electrode arrays (64 channels each) in one macaque (male, 7 years old) upon presentation with images (resized to 500 *×* 500 px) in a passive fixation experiment (Figure 4). During the experiment, 4000 training images were each presented once, which ensured that these training set responses covered a diverse set of stimulus variations (note that repetitions would limit the total number of distinct images presented). In contrast, 100/200 test set images were each presented twenty times to increase the signal-to-noise ratio, which facilitated more reliable assessment and interpretation. The images were presented in a randomized order. Next, neural responses were recorded in V1 (7 arrays), V4 (4 arrays) and IT (4 arrays) leading to a total of 960 channels (see electrode placings in Figure 2). For each trial, we averaged the early response of each channel using the following time windows: 25-125 ms for V1, 50-150 ms for V4 and 75-175 ms for IT. To capture feedforward processing in each region, the time windows were centered on the response peaks and averaged across trials and channels, as determined on an independent dataset of responses to 22k natural images. The 100 ms window length accounted for the variability of response latency across channels and stimuli. Normalization was carried out as in [76], such that for each channel, the mean response was subtracted from all the values which were then divided by the standard deviation.

To determine the contribution of the activity in each brain region to the overall model performance, we evaluated the decoder using partially occluded test set data. Concretely, we used our main decoder which was trained on neural data from all three brain areas and evaluated it using test set recordings from one brain area. To do this, the responses from the other two areas were occluded by the average response of all but the corresponding response. Alternatively, one could also evaluate the contribution per region by training three *independent* decoders on subsets of neural data (V1-only, V4-only and IT-only) which would allow for evaluation of the contribution of each brain area independently of one another. But in our case, we used the occlusion approach to investigate the area-specific contribution to the *same* decoder’s performance by keeping the contributions from the other two areas constant.

### 4.4 Models

We used linear mapping to evaluate our claim that the feature- and neural representation effectively encode the same stimulus properties, as is standard in neural coding [77, 6]. A more complex nonlinear transformation would not be valid to support this claim since nonlinearities will fundamentally change the underlying representations.

#### 4.4.1 Encoding

Kernel ridge regression was used to model how every recording site in the visual cortex is linearly dependent on the stimulus features. That is, an encoding model is defined for each electrode. Encoding required regularization to avoid overfitting since we predicted from feature space **x***_i_ → ϕ*(**x***_i_*) where *ϕ*() is the feature extraction model. Hence we used ridge regression where the norm of **w** is penalized to define encoding models by a weighted sum of *ϕ*(**x_i_**):

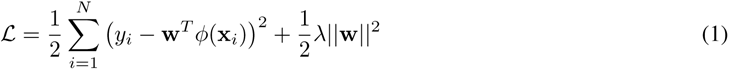

where **x** = (**x**_1_, **x**_2_*, . . . ,* **x***_N_* )*^T^ ∈* R*^N×d^*, *y* = (*y*_1_*, y*_2_*, . . . , y_N_* )*^T^ ∈* R*^N×^*^1^, *N* the number of stimulus-response pairs, *d* the number of pixels, and *λ ≥* 0 the regularization parameter. We then solved for **w** by applying the “kernel trick” [78]:

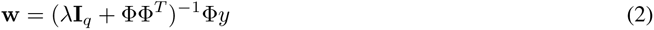

where Φ = (*ϕ*(*x*_1_)*, ϕ*(*x*_2_)*, . . . , ϕ*(*x_N_* ))*^T^ ∈* R*^N×q^* (i.e., the design matrix) where *q* is the number of feature elements and *y* = (*y*_1_*, y*_2_*, . . . , y_N_* )*^T^ ∈* R*^N×^*^1^. This means that **w** must lie in the space induced by the training data even when *q ≫ N* . The optimal *λ* is determined with grid search, as in [2]. The grid is obtained by dividing the domain of *λ* in *M* values and evaluating model performance at every value. This hyperparameter domain is controlled by the capacity of the model, i.e., the effective degrees of freedom dof of the ridge regression fit from [1*, N* ]:

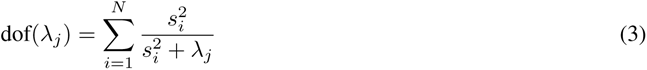

where *s* are the non-zero singular values of the design matrix Φ as obtained by singular value decomposition. We can solve for each *λ_j_* with Newton’s method. Now that the grid of lambda values is defined, we can search for the optimal *λ_j_* that minimizes the 5-fold cross-validation error.

#### 4.4.2 Decoding

Multiple linear regression was used to model how the individual units within feature representations *y_i_* (e.g., *w_i_*-latents) are linearly dependent on brain activity **x_i_**:

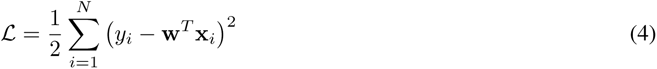

where *i* ranges over samples. We reconstructed the images by feeding the predicted latents to brain responses of the test set by feeding them to the generator without truncation.

### 4.5 Evaluation

Decoding performance was evaluated by six metrics that compared the stimuli from the held-out test set with their reconstructions from brain activity: perceptual cosine similarity using the five MaxPool layer outputs of VGG16 and latent cosine similarity. For *perceptual cosine similarity*, we computed the cosine similarity between layer activations (rather than pixel space which is the model input) extracted by VGG16 pretrained for object recognition. This metric reflects human perception of similarity better because it takes more high-level visual cues into account (e.g., color, texture, and spatial information) and human perception is often not directly related to the pixel values themselves. Specifically, we fed the stimuli and their reconstructions to the DNN and then considered the cosine similarity per activation unit:

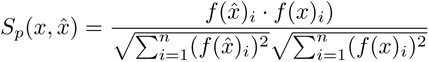

where *x* and *x*^ are the visual stimuli and their reconstructions, respectively, *n* the number of activation elements, and *f* (.) the image-activation transformation. For *latent similarity*, we considered the cosine similarity per latent dimension between predicted and ground-truth latent vectors:

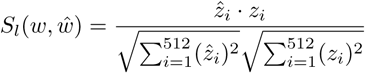

where *w*^ and *w* are the 512-dimensional predicted and ground-truth feature-disentangled latent vectors, respectively.

### 4.6 Implementation details

All analyses were carried out in Python 3.8 on a cloud-based virtual machine with Intel(R) Xeon(R) CPU @ 2.20GHz and NVIDIA Tesla T4 GPU (Driver Version: 510.47.03, CUDA Version: 11.6) on a Linux-based operating system. We used the original PyTorch implementations of StyleGAN3 and StyleGAN-XL to generate the faces and natural images in this manuscript. We used VGG16 for face recognition and object recognition for analysis of the faces and natural images. The scripts to generate the visual datasets as well as our implementations of neural encoding and -decoding can be found on our GitHub repository.

### 4.7 Ethics statement

All procedures complied with the NIH Guide for Care and Use of Laboratory Animals and were approved by the local institutional animal care and use committee of the Royal Netherlands Academy of Arts and Sciences.

In conjunction with the evolving field of neural decoding grows the concern regarding mental privacy [79] — a concept that safeguards the sanctity of individual cognitive experiences. Importantly, our methodology included extensive datasets for which constant and complete subject cooperation was required throughout the process to decode very specific information from the brain. Together with the invasive nature of our approach, which entails surgical interventions, this presents substantial barriers to any unsolicited invasion of mental privacy. Furthermore, it is important to at all times strictly follow ethical rules and regulations that govern data extraction, storage, and protection. Finally, this work solely concentrated on reconstructing visual perception; it has not extended into the domains of imagery or dreams which are more closely aligned with private cognitive experiences.

## Acknowledgements

We thank Kor Brandsma, Anneke Ditewig, Taijsha van Rees and Lex Beekman for biotechnical support.

## List of captions from supplementary information

### S1 Results for macaque #2

- **S1.1 Encoding performance.** The effectiveness of each encoding model is assessed using the Pearson correlation coefficients between predicted and recorded neural responses. The first and second graph denote discriminative and generative representations, respectively.
- **S1.2 Generative-based encoding performance.** For each individual microelectrode unit, we fit three encoding models based on three distinct feature representations: *z*-, *w−* and CLIP-latent representations. As such, we fit 3*×* 1020 independent encoders, resulting in 3*×* 1024 predicted neural responses. The scatterplots display the prediction-target correlation (*r*) of one encoding model on the X-axis and another encoding model on the Y-axis to investigate the relationship between the two. Each dot represents the performance of one modeled microelectrode unit in terms of both encoding models (so, 1024 dots per plot). The diagonal represents equal performance between both models. It is clear to see that *w*-latents always outperform *z*- and CLIP-latents because most dots lie in the direction of the *w*-axis (above the diagonal).
- **S1.3 Qualitative results.** This figure shows the 100 test set stimuli (top row) and their reconstructions from brain activity from subject 1 (middle row) and subject 2 (bottom row).

### S2 Reconstruction via ***z***-latents

- **S2.1 Qualitative results for face images:** test set stimuli (top), ’original’ reconstructions from brain activity via *w*-latents (middle) and reconstructions from brain activity via *z*-latents.

- **S2.2 Qualitative results for natural images:** test set stimuli (top), ’original’ reconstructions from brain activity via *w*-latents (middle) and reconstructions from brain activity via *z*-latents.

### S3 Reconstruction baseline

- **S3.1 Quantitative results.** Reconstruction performance (*mean ± std.error*) in terms of six metrics of perceptual cosine similarity using the five MaxPool layer outputs of VGG16 for face or image recognition and latent cosine similarity between *w*-latents of stimuli and their reconstructions when using the recordings from all recording sites (i.e., V1, V4 and IT together). The first row shows the original reconstruction performance from the manuscript, and the second and third row of the baseline using the prior of 10,000 and 6,000,000 images, respectively.
- **S3.2 Qualitative results for face images (prior=10,000):** test set stimuli (top), ’original’ reconstructions from brain activity using linear decoding (middle) and reconstructions from brain activity using the baseline approach.
- **S3.3 Qualitative results for face images (prior=6,000,000):** test set stimuli (top), ’original’ reconstructions from brain activity using linear decoding (middle) and reconstructions from brain activity using the baseline approach.
- **S3.3 Qualitative results for natural images (prior=10,000):** test set stimuli (top), ’original’ reconstructions from brain activity using linear decoding (middle) and reconstructions from brain activity using the baseline approach.
- **S3.4 Qualitative results for natural images (prior=60,000,000):** test set stimuli (top), ’original’ reconstructions from brain activity using linear decoding (middle) and reconstructions from brain activity using the baseline approach.

### S4 Leave-one-example-out analysis

- **4.1 Quantitative results.** Reconstruction performance (*mean ± std.error*) in terms of six metrics of perceptual cosine similarity using the five MaxPool layer outputs of VGG16 for object recognition and latent cosine similarity between *w*-latents of stimuli and their reconstructions when using the recordings from all recording sites (i.e., V1, V4 and IT together). The first row shows the original reconstruction performance from the manuscript and the second row of the leave-one-example-out analysis.
- **S4.2 Qualitative reconstruction results:** training examples that are used for testing (top) row and their reconstructions from brain activity (bottom row) via *w*-latents.

### S5 Leave-one-class-out analysis

- **5.1 Quantitative results.** Reconstruction performance (*mean ± std.error*) in terms of six metrics of perceptual cosine similarity using the five MaxPool layer outputs of VGG16 for object recognition and latent cosine similarity between *w*-latents of stimuli and their reconstructions when using the recordings from all recording sites (i.e., V1, V4 and IT together). The first row shows the original reconstruction performance from the manuscript and the second row of the leave-one-class-out analysis.
- **5.2 Qualitative reconstruction results:** test set stimuli (top) and their reconstructions from brain activity when the training examples of their class are excluded from training (middle). The original reconstructions, when all classes are included during training, are also displayed for reference.

### S6 Permutation test analysis

- **6.1 Permutation test analysis.** The quantitative results were verified with a permutation test as follows: per iteration, 100 and 200 latents (and their corresponding images) were randomly sampled for faces and natural images, respectively, to evaluate their similarity to the stimuli in terms of the six similarity metrics. In the above graphs, these similarity metrics were plotted over 100 iterations and we discovered that random samples were never better than our predictions from brain activity.

### S7 Visual guide

- **7.1 Visual guide.** For the six similarity metrics, we display the 5 lowest and highest stimulus-reconstruction pairs from the dataset of faces- (left) and natural images dataset (right). The top row denotes the stimulus and the bottom row the reconstruction from brain activity.

### S8 Generating abstract images

- **8.1 Generating abstract images.** Top: abstract image (taken from Shen et al. (2019)). Bottom: image corresponding to the iteratively-optimized latent to match its visual features with those of the target latent.

### S9 Category labels (Tiny ImageNet [73])

## References

[1] Winrich A Freiwald and Doris Y Tsao. Functional compartmentalization and viewpoint generalization within the macaque face-processing system. Science, 330(6005):845–851, 2010.

[2] Umut Güçlü and Marcel van Gerven. Unsupervised feature learning improves prediction of human brain activity in response to natural images. PLoS computational biology, 10(8):e1003724, 2014.

[3] Daniel LK Yamins, Ha Hong, Charles F Cadieu, Ethan A Solomon, Darren Seibert, and James J DiCarlo. Performance-optimized hierarchical models predict neural responses in higher visual cortex. Proceedings of the national academy of sciences, 111(23):8619–8624, 2014.

[4] Charles F Cadieu, Ha Hong, Daniel LK Yamins, Nicolas Pinto, Diego Ardila, Ethan A Solomon, Najib J Majaj, and James J DiCarlo. Deep neural networks rival the representation of primate it cortex for core visual object recognition. PLoS computational biology, 10(12):e1003963, 2014.

[5] Seyed-Mahdi Khaligh-Razavi and Nikolaus Kriegeskorte. Deep supervised, but not unsupervised, models may explain it cortical representation. PLoS computational biology, 10(11):e1003915, 2014.

[6] Umut Güçlü and Marcel van Gerven. Deep neural networks reveal a gradient in the complexity of neural representations across the ventral stream. Journal of Neuroscience, 35(27):10005–10014, 2015.

[7] Daniel LK Yamins and James J DiCarlo. Using goal-driven deep learning models to understand sensory cortex. Nature neuroscience, 19(3):356–365, 2016.

[8] Radoslaw Martin Cichy, Aditya Khosla, Dimitrios Pantazis, Antonio Torralba, and Aude Oliva. Comparison of deep neural networks to spatio-temporal cortical dynamics of human visual object recognition reveals hierarchical correspondence. Scientific reports, 6(1):1–13, 2016.

[9] Umut Güçlü, Jordy Thielen, Michael Hanke, and Marcel van Gerven. Brains on beats. Advances in Neural Information Processing Systems, 29, 2016.

[10] Marcel van Gerven. A primer on encoding models in sensory neuroscience. Journal of Mathematical Psychology, 76:172–183, 2017.

[11] Michael Eickenberg, Alexandre Gramfort, Gaël Varoquaux, and Bertrand Thirion. Seeing it all: Convolutional network layers map the function of the human visual system. NeuroImage, 152:184–194, 2017.

[12] Le Chang and Doris Y Tsao. The code for facial identity in the primate brain. Cell, 169(6):1013–1028, 2017.

[13] Umut Güçlü and Marcel van Gerven. Probing human brain function with artificial neural networks. Computational Models of Brain and Behavior, pages 413–423, 2017.

[14] Katja Seeliger, Matthias Fritsche, Umut Güçlü, Sanne Schoenmakers, J Schoffelen, Sander Bosch, and M van Gerven. Convolutional neural network-based encoding and decoding of visual object recognition in space and time. NeuroImage, 180:253–266, 2018.

[15] James V Haxby, M Ida Gobbini, Maura L Furey, Alumit Ishai, Jennifer L Schouten, and Pietro Pietrini. Distributed and overlapping representations of faces and objects in ventral temporal cortex. Science, 293(5539):2425–2430, 2001.

[16] Yukiyasu Kamitani and Frank Tong. Decoding the visual and subjective contents of the human brain. Nature neuroscience, 8(5):679–685, 2005.

[17] Dustin E Stansbury, Thomas Naselaris, and Jack L Gallant. Natural scene statistics account for the representation of scene categories in human visual cortex. Neuron, 79(5):1025–1034, 2013.

[18] Alexander G Huth, Tyler Lee, Shinji Nishimoto, Natalia Y Bilenko, An T Vu, and Jack L Gallant. Decoding the semantic content of natural movies from human brain activity. Frontiers in systems neuroscience, 10:81, 2016.

[19] Tomoyasu Horikawa and Yukiyasu Kamitani. Generic decoding of seen and imagined objects using hierarchical visual features. Nature communications, 8(1):1–15, 2017.

[20] Tom M Mitchell, Svetlana V Shinkareva, Andrew Carlson, Kai-Min Chang, Vicente L Malave, Robert A Mason, and Marcel Adam Just. Predicting human brain activity associated with the meanings of nouns. Science, 320(5880):1191–1195, 2008.

[21] Kendrick N Kay, Thomas Naselaris, Ryan J Prenger, and Jack L Gallant. Identifying natural images from human brain activity. Nature, 452(7185):352–355, 2008.

[22] Umut Güçlü and Marcel van Gerven. Increasingly complex representations of natural movies across the dorsal stream are shared between subjects. NeuroImage, 145:329–336, 2017.

[23] Umut Güçlü and Marcel van Gerven. Modeling the dynamics of human brain activity with recurrent neural networks. Frontiers in computational neuroscience, 11:7, 2017.

[24] Bertrand Thirion, Edouard Duchesnay, Edward Hubbard, Jessica Dubois, Jean-Baptiste Poline, Denis Lebihan, and Stanislas Dehaene. Inverse retinotopy: inferring the visual content of images from brain activation patterns. NeuroImage, 33(4):1104–1116, 2006.

[25] Yoichi Miyawaki, Hajime Uchida, Okito Yamashita, Masa-aki Sato, Yusuke Morito, Hiroki C Tanabe, Norihiro Sadato, and Yukiyasu Kamitani. Visual image reconstruction from human brain activity using a combination of multiscale local image decoders. Neuron, 60(5):915–929, 2008.

[26] Thomas Naselaris, Ryan J Prenger, Kendrick N Kay, Michael Oliver, and Jack L Gallant. Bayesian reconstruction of natural images from human brain activity. Neuron, 63(6):902–915, 2009.

[27] Marcel van Gerven, Floris P de Lange, and Tom Heskes. Neural decoding with hierarchical generative models. Neural computation, 22(12):3127–3142, 2010.

[28] Shinji Nishimoto, An T Vu, Thomas Naselaris, Yuval Benjamini, Bin Yu, and Jack L Gallant. Reconstructing visual experiences from brain activity evoked by natural movies. Current biology, 21(19):1641–1646, 2011.

[29] Sanne Schoenmakers, Markus Barth, Tom Heskes, and Marcel Van Gerven. Linear reconstruction of perceived images from human brain activity. NeuroImage, 83:951–961, 2013.

[30] Umut Güçlü and Marcel van Gerven. Unsupervised learning of features for bayesian decoding in functional magnetic resonance imaging. In Belgian-Dutch Conference on Machine Learning, 2013.

[31] Alan S Cowen, Marvin M Chun, and Brice A Kuhl. Neural portraits of perception: reconstructing face images from evoked brain activity. NeuroImage, 94:12–22, 2014.

[32] Changde Du, Changying Du, and Huiguang He. Sharing deep generative representation for perceived image reconstruction from human brain activity. In 2017 International Joint Conference on Neural Networks (IJCNN), pages 1049–1056. IEEE, 2017.

[33] Yağmur Güçlütürk, Umut Güçlü, Katja Seeliger, Sander Bosch, Rob van Lier, and Marcel van Gerven. Re-constructing perceived faces from brain activations with deep adversarial neural decoding. Advances in neural information processing systems, 30, 2017.

[34] Guohua Shen, Tomoyasu Horikawa, Kei Majima, and Yukiyasu Kamitani. Deep image reconstruction from human brain activity. PLoS computational biology, 15(1):e1006633, 2019.

[35] Rufin VanRullen and Leila Reddy. Reconstructing faces from fmri patterns using deep generative neural networks. Communications biology, 2(1):1–10, 2019.

[36] Thirza Dado, Yağmur Güçlütürk, Luca Ambrogioni, Gabriëlle Ras, Sander Bosch, Marcel van Gerven, and Umut Güçlü. Hyperrealistic neural decoding for reconstructing faces from fmri activations via the gan latent space. Scientific reports, 12(1):1–9, 2022.

[37] Nadine Dijkstra, Sander Bosch, and Marcel van Gerven. Shared neural mechanisms of visual perception and imagery. Trends in Cognitive Sciences, 23(5):423–434, 2019.

[38] Ian Goodfellow, Jean Pouget-Abadie, Mehdi Mirza, Bing Xu, David Warde-Farley, Sherjil Ozair, Aaron Courville, and Yoshua Bengio. Generative adversarial nets. Advances in neural information processing systems, 27, 2014.

[39] Andrew Brock, Jeff Donahue, and Karen Simonyan. Large scale gan training for high fidelity natural image synthesis. arXiv preprint arXiv:1809.11096, 2018.

[40] Tero Karras, Timo Aila, Samuli Laine, and Jaakko Lehtinen. Progressive growing of gans for improved quality, stability, and variation. arXiv preprint arXiv:1710.10196, 2017.

[41] Tero Karras, Samuli Laine, and Timo Aila. A style-based generator architecture for generative adversarial networks. In Proceedings of the IEEE/CVF Conference on Computer Vision and Pattern Recognition, pages 4401–4410, 2019.

[42] Tero Karras, Miika Aittala, Samuli Laine, Erik Härkönen, Janne Hellsten, Jaakko Lehtinen, and Timo Aila. Alias-free generative adversarial networks. Advances in Neural Information Processing Systems, 34, 2021.

[43] Nikolaus Kriegeskorte. Deep neural networks: a new framework for modeling biological vision and brain information processing. Annual review of vision science, 1:417–446, 2015.

[44] Alan Yuille and Daniel Kersten. Vision as bayesian inference: analysis by synthesis? Trends in cognitive sciences, 10(7):301–308, 2006.

[45] Yujun Shen, Jinjin Gu, Xiaoou Tang, and Bolei Zhou. Interpreting the latent space of gans for semantic face editing. In Proceedings of the IEEE/CVF conference on computer vision and pattern recognition, pages 9243–9252, 2020.

[46] Irina Higgins, David Amos, David Pfau, Sebastien Racaniere, Loic Matthey, Danilo Rezende, and Alexander Lerchner. Towards a definition of disentangled representations. arXiv preprint arXiv:1812.02230, 2018.

[47] Hans Super and Pieter R Roelfsema. Chronic multiunit recordings in behaving animals: advantages and limitations. Progress in brain research, 147:263–282, 2005.

[48] Alec Radford, Jong Wook Kim, Chris Hallacy, Aditya Ramesh, Gabriel Goh, Sandhini Agarwal, Girish Sastry, Amanda Askell, Pamela Mishkin, Jack Clark, et al. Learning transferable visual models from natural language supervision. In International conference on machine learning, pages 8748–8763. PMLR, 2021.

[49] Robin Rombach, Andreas Blattmann, Dominik Lorenz, Patrick Esser, and Björn Ommer. High-resolution image synthesis with latent diffusion models. In Proceedings of the IEEE/CVF Conference on Computer Vision and Pattern Recognition, pages 10684–10695, 2022.

[50] Adrien Doerig, Tim C Kietzmann, Emily Allen, Yihan Wu, Thomas Naselaris, Kendrick Kay, and Ian Charest. Semantic scene descriptions as an objective of human vision. arXiv preprint arXiv:2209.11737, 2022.

[51] Aria Y Wang, Kendrick Kay, Thomas Naselaris, Michael J Tarr, and Leila Wehbe. Better models of human high-level visual cortex emerge from natural language supervision with a large and diverse dataset. Nature Machine Intelligence, pages 1–12, 2023.

[52] L.G. Ungerleider and M. Mishkin. Two cortical visual systems. In Analysis of visual behavior, pages 549–586–. MIT Press, Cambridge, MA, 1982.

[53] David H Hubel and Torsten N Wiesel. Receptive fields, binocular interaction and functional architecture in the cat’s visual cortex. The Journal of physiology, 160(1):106–154, 1962.

[54] Charles G Gross, CE de Rocha-Miranda, and DB Bender. Visual properties of neurons in inferotemporal cortex of the macaque. Journal of neurophysiology, 35(1):96–111, 1972.

[55] Chou P Hung, Gabriel Kreiman, Tomaso Poggio, and James J DiCarlo. Fast readout of object identity from macaque inferior temporal cortex. Science, 310(5749):863–866, 2005.

[56] Martin I Sereno, AM Dale, JB Reppas, KK Kwong, JW Belliveau, TJ Brady, BR Rosen, and RBH Tootell. Borders of multiple visual areas in humans revealed by functional magnetic resonance imaging. Science, 268(5212):889–893, 1995.

[57] Mark D Lescroart and Jack L Gallant. Human scene-selective areas represent 3d configurations of surfaces. Neuron, 101(1):178–192, 2019.

[58] Tomoyasu Horikawa and Yukiyasu Kamitani. Hierarchical neural representation of dreamed objects revealed by brain decoding with deep neural network features. Frontiers in computational neuroscience, 11:4, 2017.

[59] Ghislain St-Yves and Thomas Naselaris. Generative adversarial networks conditioned on brain activity reconstruct seen images. In 2018 IEEE international conference on systems, man, and cybernetics (SMC), pages 1054–1061. IEEE, 2018.

[60] Guohua Shen, Kshitij Dwivedi, Kei Majima, Tomoyasu Horikawa, and Yukiyasu Kamitani. End-to-end deep image reconstruction from human brain activity. Frontiers in Computational Neuroscience, page 21, 2019.

[61] Milad Mozafari, Leila Reddy, and Rufin VanRullen. Reconstructing natural scenes from fmri patterns using bigbigan. In 2020 International joint conference on neural networks (IJCNN), pages 1–8. IEEE, 2020.

[62] Guy Gaziv, Roman Beliy, Niv Granot, Assaf Hoogi, Francesca Strappini, Tal Golan, and Michal Irani. Self-supervised natural image reconstruction and large-scale semantic classification from brain activity. NeuroImage, 254:119121, 2022.

[63] Kuan Han, Haiguang Wen, Junxing Shi, Kun-Han Lu, Yizhen Zhang, Di Fu, and Zhongming Liu. Variational autoencoder: An unsupervised model for encoding and decoding fmri activity in visual cortex. NeuroImage, 198:125–136, 2019.

[64] Katja Seeliger, Umut Güçlü, Luca Ambrogioni, Yagmur Güçlütürk, and Marcel van Gerven. Generative adversarial networks for reconstructing natural images from brain activity. NeuroImage, 181:775–785, 2018.

[65] Irina Higgins, Le Chang, Victoria Langston, Demis Hassabis, Christopher Summerfield, Doris Tsao, and Matthew Botvinick. Unsupervised deep learning identifies semantic disentanglement in single inferotemporal face patch neurons. Nature communications, 12(1):6456, 2021.

[66] Hang Shao, Abhishek Kumar, and P Thomas Fletcher. The riemannian geometry of deep generative models. In Proceedings of the IEEE Conference on Computer Vision and Pattern Recognition Workshops, pages 315–323, 2018.

[67] Tomas Mikolov, Ilya Sutskever, Kai Chen, Greg S Corrado, and Jeff Dean. Distributed representations of words and phrases and their compositionality. Advances in neural information processing systems, 26, 2013.

[68] James J DiCarlo and David D Cox. Untangling invariant object recognition. Trends in cognitive sciences, 11(8):333–341, 2007.

[69] Tero Karras, Samuli Laine, Miika Aittala, Janne Hellsten, Jaakko Lehtinen, and Timo Aila. Analyzing and improving the image quality of stylegan. In Proceedings of the IEEE/CVF Conference on Computer Vision and Pattern Recognition, pages 8110–8119, 2020.

[70] Xun Huang and Serge Belongie. Arbitrary style transfer in real-time with adaptive instance normalization. In Proceedings of the IEEE international conference on computer vision, pages 1501–1510, 2017.

[71] Axel Sauer, Katja Schwarz, and Andreas Geiger. Stylegan-xl: Scaling stylegan to large diverse datasets. In ACM SIGGRAPH 2022 Conference Proceedings, pages 1–10, 2022.

[72] Jia Deng, Wei Dong, Richard Socher, Li-Jia Li, Kai Li, and Li Fei-Fei. Imagenet: A large-scale hierarchical image database. In 2009 IEEE conference on computer vision and pattern recognition, pages 248–255. IEEE, 2009.

[73] Ya Le and Xuan Yang. Tiny imagenet visual recognition challenge. CS 231N, 7(7):3, 2015.

[74] Omkar M. Parkhi, Andrea Vedaldi, and Andrew Zisserman. Deep face recognition. In Xianghua Xie, Mark W. Jones, and Gary K. L. Tam, editors, Proceedings of the British Machine Vision Conference (BMVC), pages 41.1–41.12. BMVA Press, September 2015.

[75] Karen Simonyan and Andrew Zisserman. Very deep convolutional networks for large-scale image recognition. arXiv preprint arXiv:1409.1556, 2014.

[76] Pouya Bashivan, Kohitij Kar, and James J DiCarlo. Neural population control via deep image synthesis. Science, 364(6439), 2019.

[77] Thomas Naselaris, Kendrick N Kay, Shinji Nishimoto, and Jack L Gallant. Encoding and decoding in fmri. NeuroImage, 56(2):400–410, 2011.

[78] Max Welling. Kernel ridge regression. Max Welling’s classnotes in machine learning, pages 1–3, 2013.

[79] Marcello Ienca, Pim Haselager, and Ezekiel J Emanuel. Brain leaks and consumer neurotechnology. Nature biotechnology, 36(9):805–810, 2018.

