## Supplementary Materials for "Brain2GAN: Feature-disentangled neural encoding and decoding of visual perception in the primate brain"

[1] Donders Institute for Brain, Cognition and Behaviour, Radboud University, Nijmegen, Netherlands. [2] Department of Vision and Cognition, Netherlands Institute for Neuroscience, Amsterdam, Netherlands. [3] Laboratory of Visual Brain Therapy, Sorbonne University, Paris, France. [4] Department of Integrative Neurophysiology, VU Amsterdam, Amsterdam, Netherlands. [5] Department of Psychiatry, Amsterdam UMC, Amsterdam, Netherlands.

### S1 Results for macaque #2

We repeated the passive fixation experiment using brain responses from V1, V2, V3 and V4 in a second macaque (male, 9 years old) with silicone-based electrodes. Note that this subject has no electrode arrays implanted in higher-level visual areas. All set-up and collection procedures, including recording equipment and the preprocessing, were identical to those of the first macaque. The encoding performance was quantified as the Pearson product-moment correlation coefficient between the predicted and recorded responses (Figure 2). Among the generative-based encoders, the $w$-latent-based encoder significantly outperformed the $z$-based encoder (2-Sample T-Test; $t(2046) = -8.3180$, $p = 1.6ee - 16$) but not the CLIP-based encoder (2-Sample T-Test; $t(2046) = -0.7301$, $p = 0.4653$). In addition, CLIP-based encoding also outperformed $z$-based encoding (2-Sample T-Test; $t(2046) = 7.6463$, $p = 3.16\text{e}{-14}$).

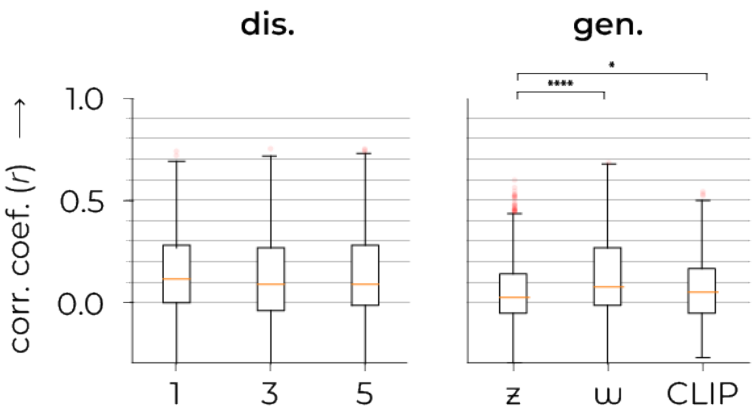


Figure 1: **S1.1 Encoding performance.** The effectiveness of each encoding model is assessed using the Pearson correlation coefficients between predicted and recorded neural responses. The first and second graph denote discriminative and generative representations, respectively.

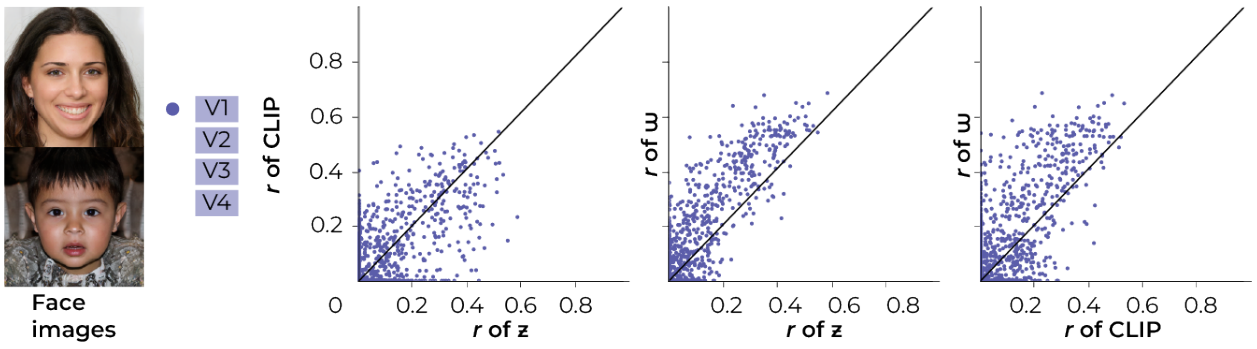


Figure 2: **S1.2 Generative-based encoding performance.** For each individual microelectrode unit, we fit three encoding models based on three distinct feature representations: $z$-, $w-$ and CLIP-latent representations. As such, we fit $3\times$ 1020 independent encoders, resulting in $3\times$ 1024 predicted neural responses. The scatterplots display the prediction-target correlation ($r$) of one encoding model on the X-axis and another encoding model on the Y-axis to investigate the relationship between the two. Each dot represents the performance of one modeled microelectrode unit in terms of both encoding models (so, 1024 dots per plot). The diagonal represents equal performance between both models. It is clear to see that $w$-latents always outperform $z$- and CLIP-latents because most dots lie in the direction of the $w$-axis (above the diagonal).

The reconstruction from areas V1, V2, V3 and V4 in the second macaque ((Figure 3, bottom row) were highly similar to the perceived stimuli but contained consistently less intricate detail compared to the reconstructions from areas V1, V4 and IT in the first macaque (Figure 3, middle row - for reference).

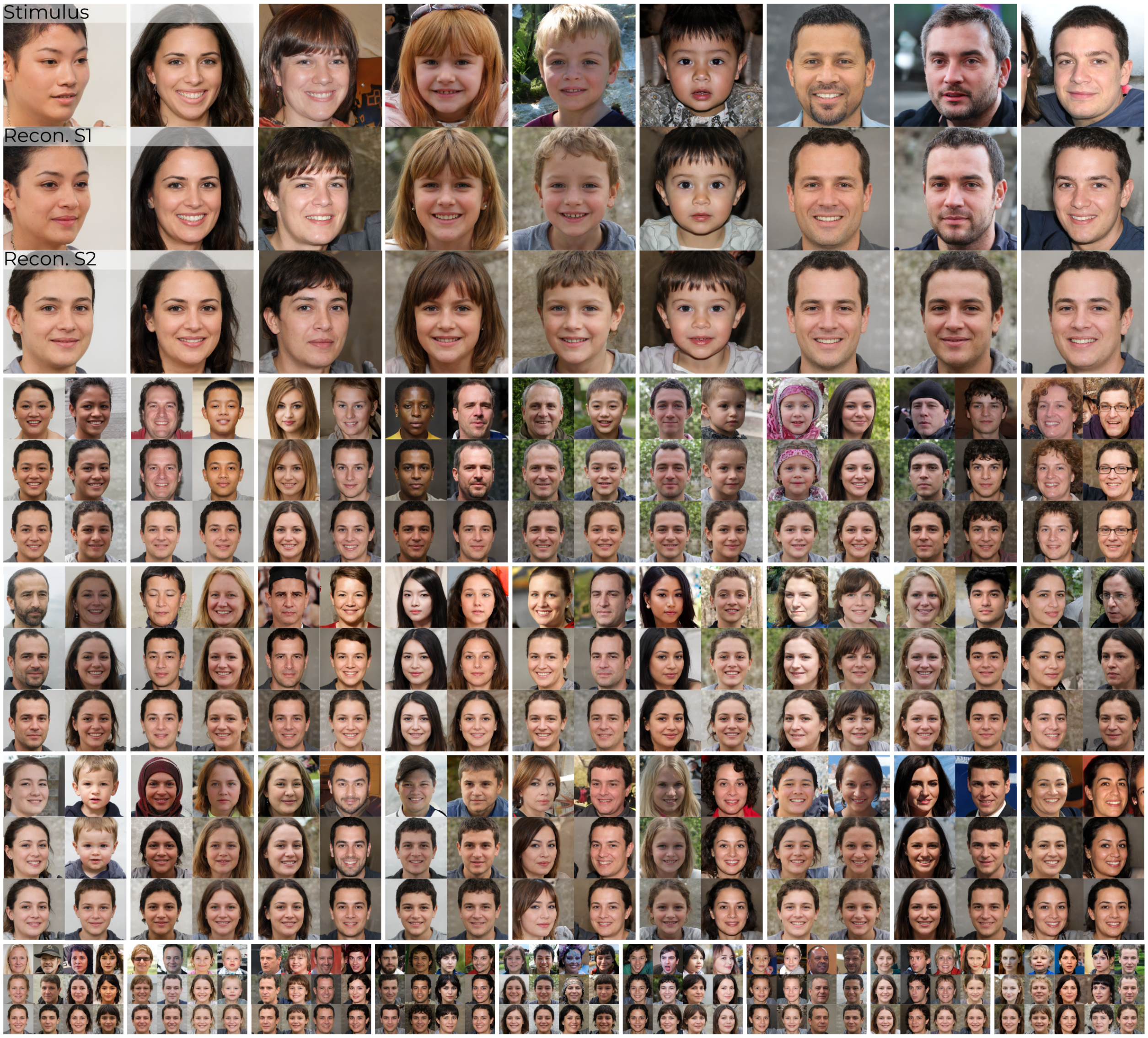


Figure 3: **S1.3 Qualitative results.** This figure shows the 100 test set stimuli (top row) and their reconstructions from brain activity in V1, V4 and IT from subject 1 (middle row) and subject 2 (bottom row).

#### S2 Reconstruction via $z$-latents

The reconstructions from $z$-latents not only demonstrate superior performance using $w$-latents in *conditional* image generation (Figure 4) but also that this disentanglement enables *unconditional* image generation using GANs (Figure 5).

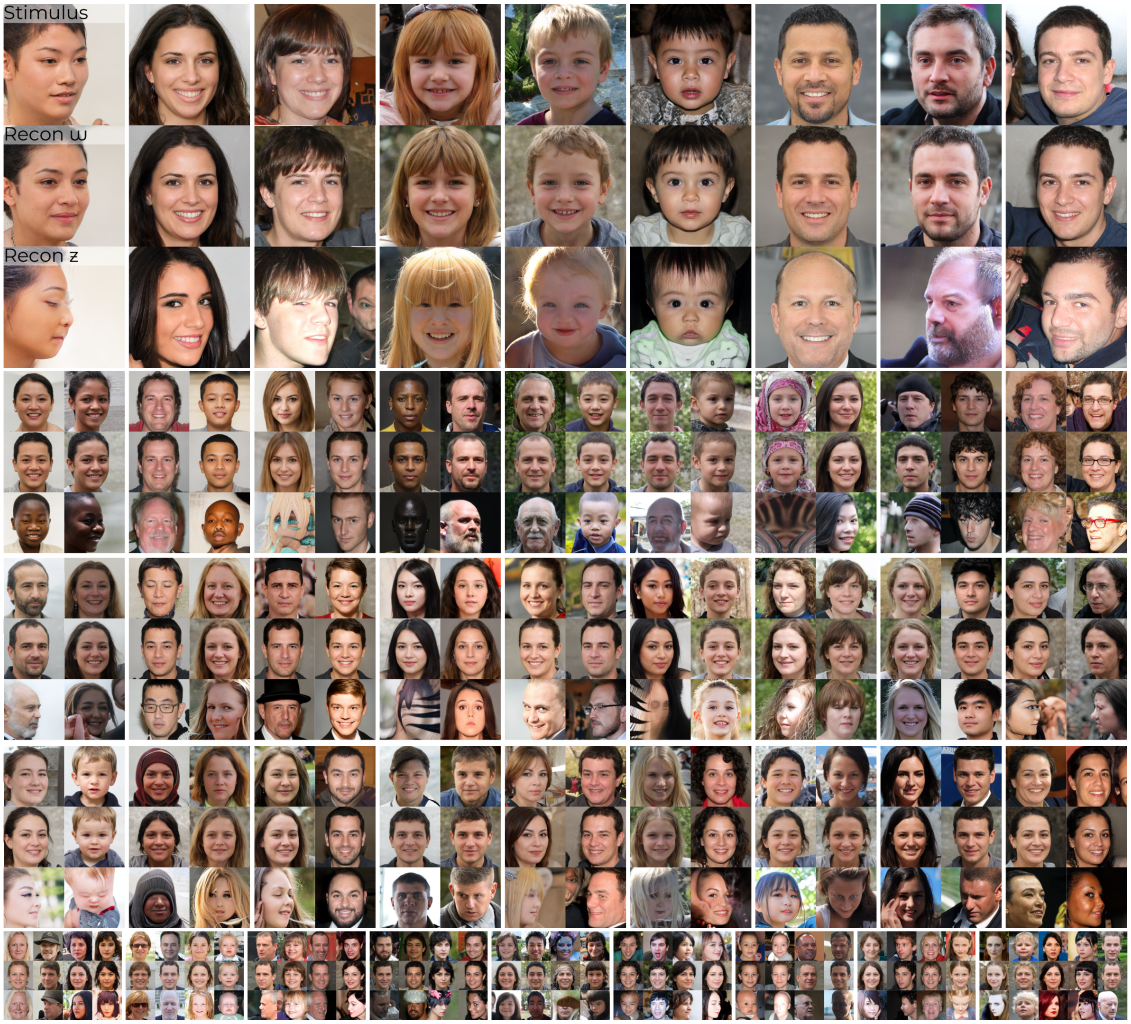


Figure 4: **S2.1 Qualitative results for face images:** test set stimuli (top), 'original' reconstructions from brain activity in V1, V4 and IT via $w$-latents (middle) and reconstructions from brain activity in V1, V4 and IT via $z$-latents.

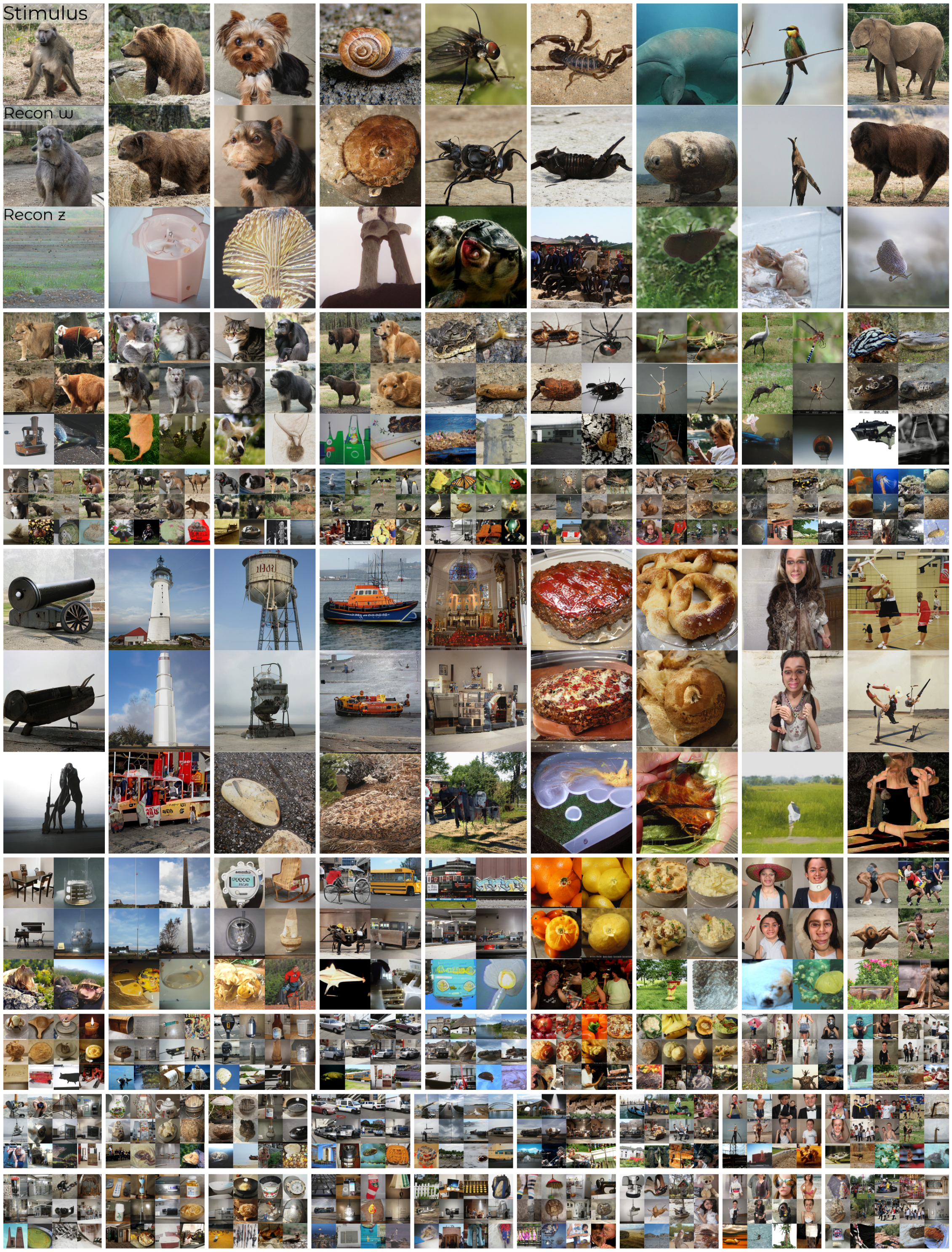


Figure 5: **S2.2 Qualitative results for natural images:** test set stimuli (top), 'original' reconstructions from brain activity in V1, V4 and IT via $w$-latents (middle) and reconstructions from brain activity in V1, V4 and IT via $z$-latents.

#### S3 Reconstruction baseline

We implemented a reconstruction method based on the work of [1] which estimated the posterior probability of a stimulus given the target responses, considering both the likelihood and prior probability of that stimulus. We modified this approach to incorporate the advantages of generative modeling, thereby aligning it with our central theme and enhancing reconstruction quality. In brief, we sampled a large number ($N = 10,000$ and $N = 6,000,000$ [2]) of $w$-latent vectors and fed them to our encoder to predict neural responses. For each test set example, we selected a subset ($n = 100$) of responses that were most similar to the observed test set response and averaged their corresponding $w$-latents. This averaged latent was fed to the generator for reconstruction. The results for faces and natural images are displayed in Figure 6, 6 and **??**, 9, respectively. Quantitative results can be found in Table 1.

Perceptually, it is easy to see that our original reconstructions outperformed those by the baseline. To our surprise, we also observed that the reconstructions using the larger prior of 6 million images were perceptually inferior compared to those using the smaller prior of 10,000 images, despite selecting what are assumed to be the best hundred latents based on their predicted similarity to the brain responses. The performance discrepancy between our decoding method and the baseline could be due to the fundamental differences in how each method relates brain activity to latents. That is, our reconstruction method utilizes a multivariate approach from all responses to predict each latent dimension independently (512 in total) such that the combined effect of all responses is captured per latent dimension. In turn, the baseline method employs the encoding model which is a mass univariate approach that predicts single neural responses (960 in total) from all latent dimensions. The encoding performance may not directly correlate with the reconstruction performance because the encoding model does not consider how all neural responses jointly relate to each latent dimension. As such, this could result in the selection of latents that are suboptimal for the purpose of visual reconstruction, even if they are good at predicting individual neural responses.

Table 1: **S3.1 Quantitative results.** Reconstruction performance ($mean \pm std.error$) in terms of six metrics of perceptual cosine similarity using the five MaxPool layer outputs of VGG16 for face or image recognition and latent cosine similarity between $w$-latents of stimuli and their reconstructions when using the recordings from all recording sites (i.e., V1, V4 and IT together). The first row shows the original reconstruction performance from the manuscript, and the second and third row of the baseline using the prior of 10,000 and 6,000,000 images, respectively.

| | | **VGG16-1 sim.** | **VGG16-2 sim.** | **VGG16-3 sim.** | **VGG16-4 sim.** | **VGG16-5 sim.** | **Lat. sim.** |
|---|---|---|---|---|---|---|---|
| Face images | orig. | $0.7871 \pm 0.0102$ | $0.7681 \pm 0.0075$ | $0.5874 \pm 0.0075$ | $0.6170 \pm 0.0085$ | $0.5940 \pm 0.0104$ | $0.5548 \pm 0.0045$ |
| | 10k | $0.6380 \pm 0.0133$ | $0.6712 \pm 0.0059$ | $0.4628 \pm 0.0059$ | $0.4702 \pm 0.0080$ | $0.4241 \pm 0.0104$ | $0.4762 \pm 0.0055$ |
| | 6M | $0.5873 \pm 0.0118$ | $0.6214 \pm 0.0048$ | $0.3961 \pm 0.0048$ | $0.3599 \pm 0.0082$ | $0.3370 \pm 0.0131$ | $0.4068 \pm 0.0071$ |
| Natural images | orig. | $0.4083 \pm 0.0036$ | $0.3322 \pm 0.0036$ | $0.2555 \pm 0.0025$ | $0.2192 \pm 0.0043$ | $0.2497 \pm 0.0066$ | $0.8032 \pm 0.0032$ |
| | 10k | $0.3723 \pm 0.0050$ | $0.3050 \pm 0.0026$ | $0.2267 \pm 0.0026$ | $0.1636 \pm 0.0039$ | $0.1438 \pm 0.0061$ | $0.7563 \pm 0.0049$ |
| | 6M | $0.3730 \pm 0.0050$ | $0.3041 \pm 0.0024$ | $0.2218 \pm 0.0024$ | $0.1545 \pm 0.0031$ | $0.1341 \pm 0.0057$ | $0.5963 \pm 0.0099$ |

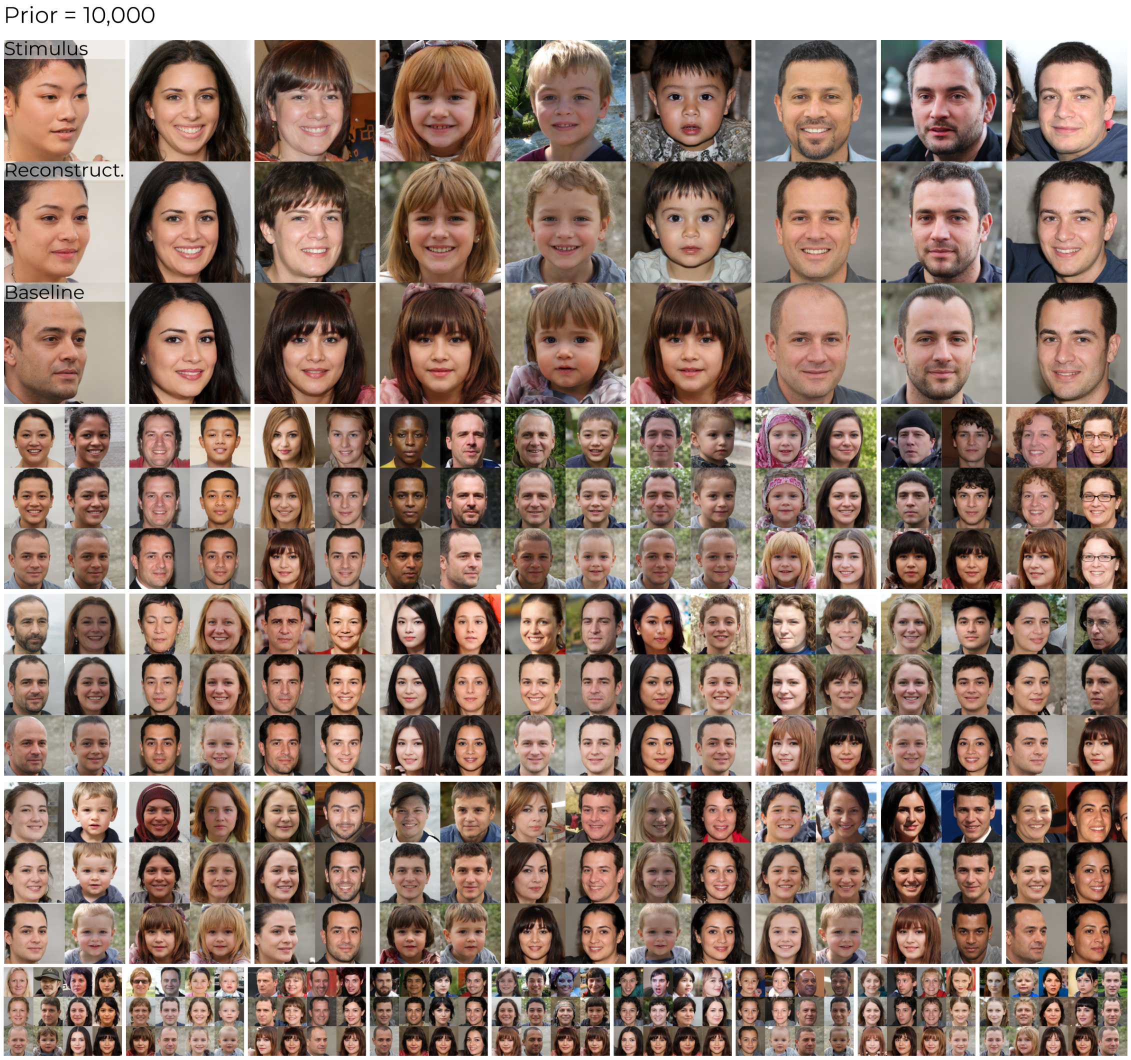


Figure 6: **S3.2 Qualitative results for face images (prior=10,000):** test set stimuli (top), 'original' reconstructions from brain activity in V1, V4 and IT using linear decoding (middle) and reconstructions from brain activity in V1, V4 and IT using the baseline approach.

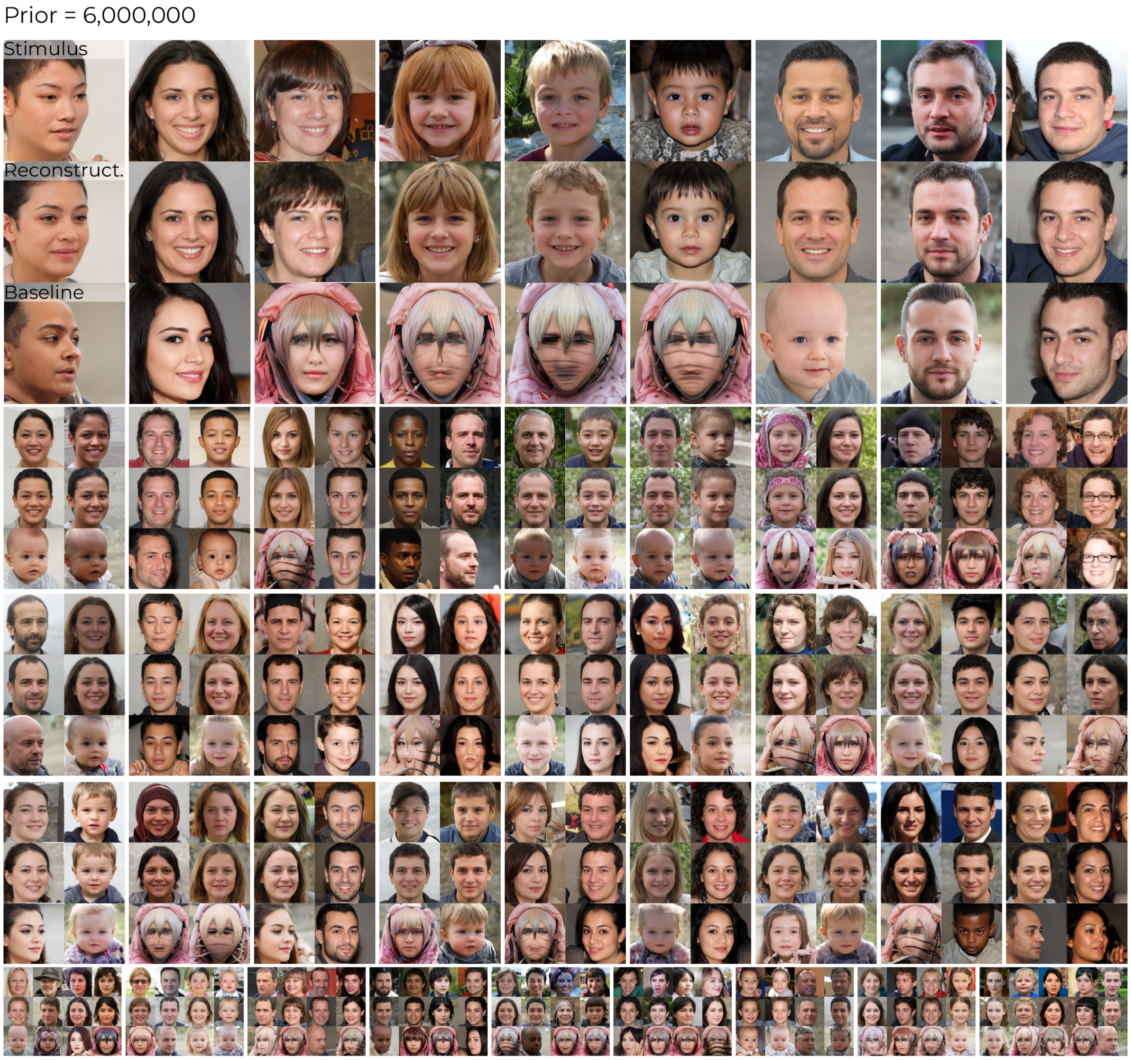


Figure 7: **S3.3 Qualitative results for face images (prior=6,000,000):** test set stimuli (top), 'original' reconstructions from brain activity in V1, V4 and IT using linear decoding (middle) and reconstructions from brain activity in V1, V4 and IT using the baseline approach.

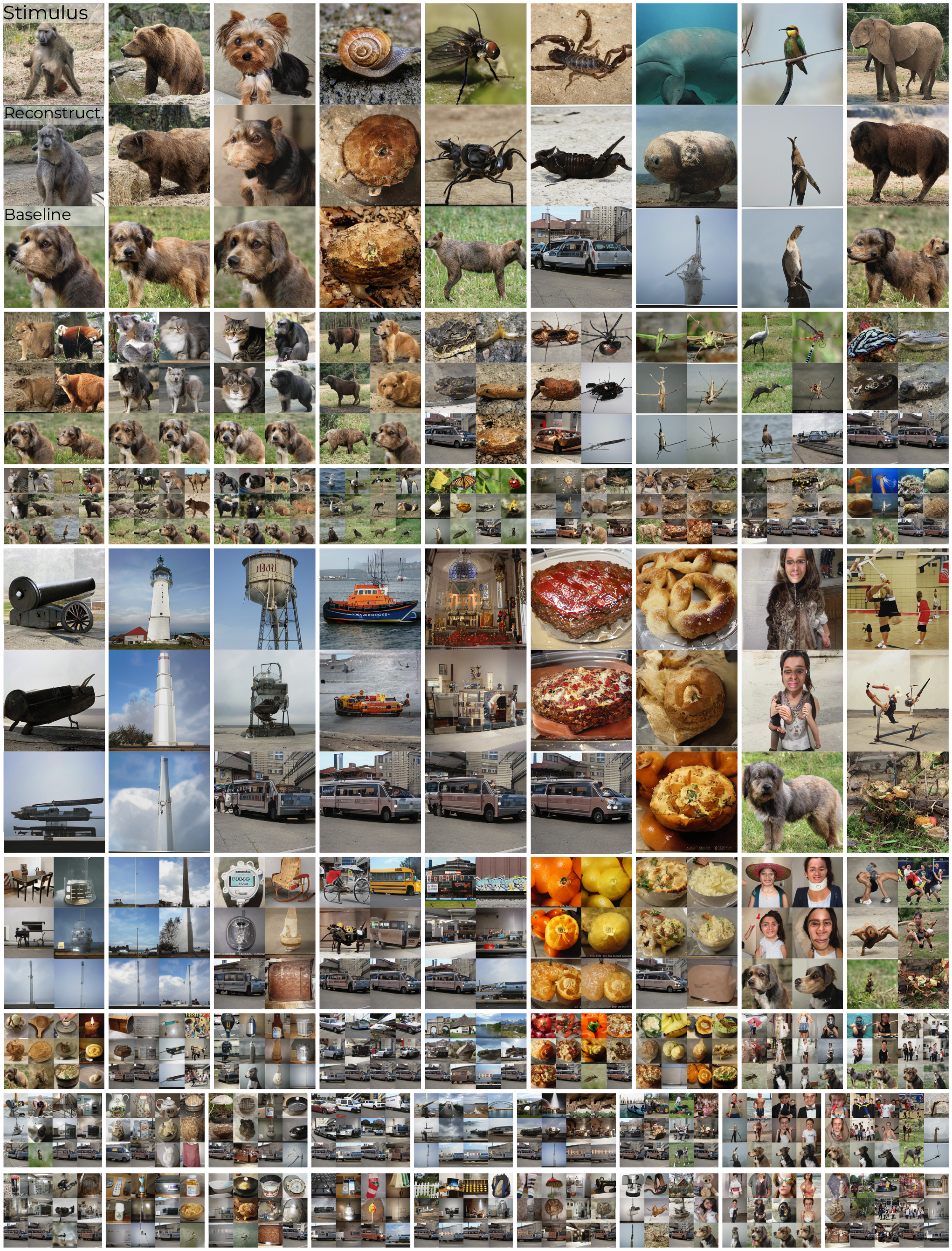


Figure 8: **S3.4 Qualitative results for natural images (prior=10,000):** test set stimuli (top), 'original' reconstructions from brain activity in V1, V4 and IT using linear decoding (middle) and reconstructions from brain activity in V1, V4 and IT using the baseline approach.

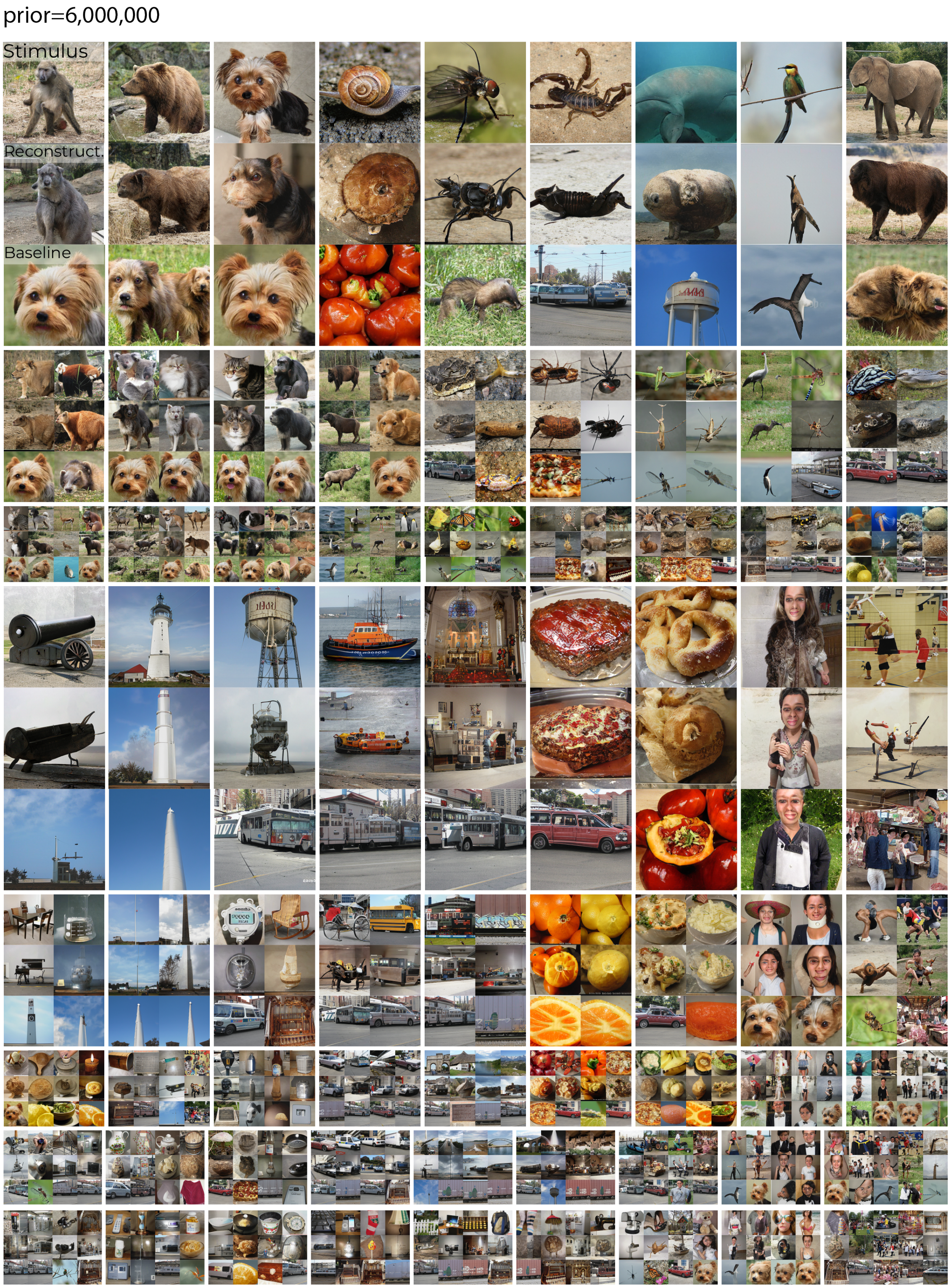


Figure 9: **S3.5 Qualitative results for natural images (prior=6,000,000):** test set stimuli (top), 'original' reconstructions from brain activity in V1, V4 and IT using linear decoding (middle) and reconstructions from brain activity in V1, V4 and IT using the baseline approach.

### S4 Leave-one-example-out analysis

In our study, we used the average $w$-latent of each class for the test set since it guaranteed high image quality and because variation was not required as only one image per category was needed. The average $w$-latent lives in the same w-space as the other $w$-latents, so there should be no advantage during reconstruction. To validate that our results were not a confound of using the average $w$-latent, we conducted an additional analysis where we trained a decoder on the first 19 training examples of each class and tested it on the remaining training example. We found that the reconstructions still resembled the stimuli, although there was a noticeable decrease in quality. This was attributed to the lower signal-to-noise ratio resulting from using a single training response. Some stimulus-reconstruction examples are presented in Figure 10A. For reference, Figure 10B shows the reconstructions based on the "original" test set. These were obtained using all training examples but decoded from a single response in the test set. This approach was chosen to match the signal-to-noise ratio conditions of the aforementioned leave-one-example-out analysis. Furthermore, Table 2 details the performance metrics of these leave-one-example-out reconstructions in comparison to the 'original' reconstruction performance based on the use of single test responses.

Table 2: **4.1 Quantitative results.** Reconstruction performance in terms of six metrics of perceptual cosine similarity using the five MaxPool layer outputs of VGG16 for object recognition and latent cosine similarity between $w$-latents of stimuli and their reconstructions ($mean \pm std.error$) when using the recordings from all recording sites (i.e., V1, V4 and IT together). The first row shows the original reconstruction performance from the manuscript and the second row of the leave-one-example-out analysis.

| | VGG16-1 sim. | VGG16-2 sim. | VGG16-3 sim. | VGG16-4 sim. | VGG16-5 sim. | Lat. sim. |
|---|---|---|---|---|---|---|
| orig. (single) | $0.3923 \pm 0.0048$ | $0.3205 \pm 0.0022$ | $0.2409 \pm 0.0022$ | $0.1870 \pm 0.0033$ | $0.1963 \pm 0.0056$ | $0.7013 \pm 0.0044$ |
| l-o-e-o. | $0.3771 \pm 0.0047$ | $0.3108 \pm 0.0022$ | $0.2317 \pm 0.0022$ | $0.1696 \pm 0.0028$ | $0.1634 \pm 0.0048$ | $0.7013 \pm 0.0044$ |

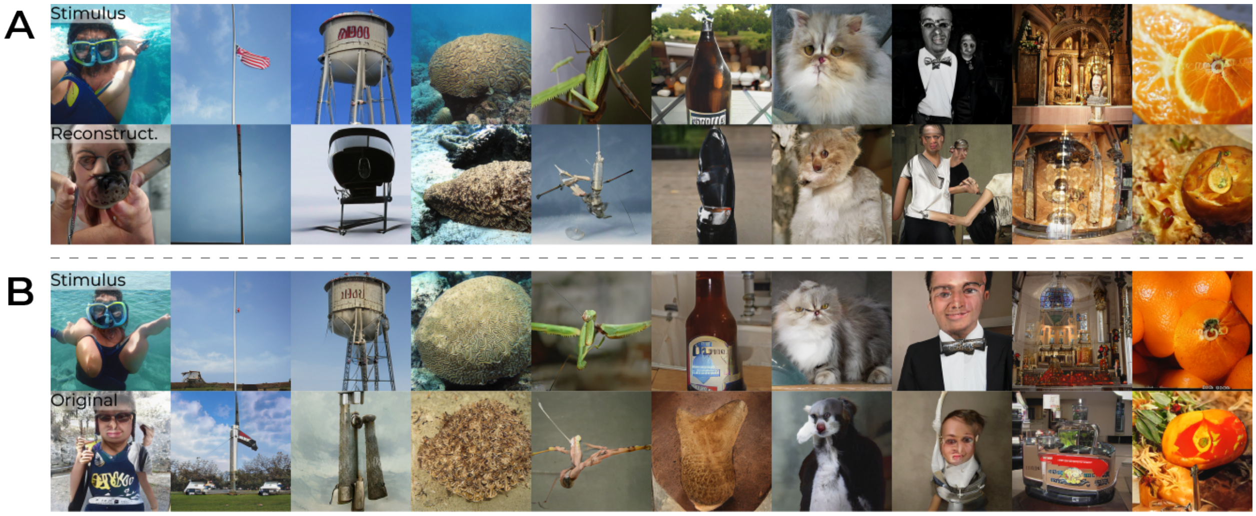


Figure 10: **S4.2 Qualitative reconstruction results:** training examples that are used for testing (top) row and their reconstructions from brain activity in V1, V4 and IT (bottom row) via $w$-latents.

### S5 Leave-one-class-out analysis

For each target category, we fit a decoder on the 199 other categories $\times$ 20 training examples, after which we predicted and reconstructed the test stimulus of the remaining class that was not included during training. We found these reconstructions to be highly consistent with the original reconstructions when all classes were included during training (Figure 11, Table 3). This finding strongly supports the assertion that our model's focus extends beyond mere classification.

Table 3: **5.1 Quantitative results.** Reconstruction performance ($mean \pm std.error$) in terms of six metrics of perceptual cosine similarity using the five MaxPool layer outputs of VGG16 for object recognition and latent cosine similarity between $w$-latents of stimuli and their reconstructions when using the recordings from all recording sites (i.e., V1, V4 and IT together). The first row shows the original reconstruction performance from the manuscript and the second row of the leave-one-class-out analysis.

| | **VGG16-1 sim.** | **VGG16-2 sim.** | **VGG16-3 sim.** | **VGG16-4 sim.** | **VGG16-5 sim.** | **Lat. sim.** |
|---|---|---|---|---|---|---|
| orig. | $0.4083 \pm 0.0036$ | $0.3322 \pm 0.0036$ | $0.2555 \pm 0.0025$ | $0.2192 \pm 0.0043$ | $0.2497 \pm 0.0066$ | $0.8032 \pm 0.0032$ |
| l-o-c-o. | $0.4039 \pm 0.0052$ | $0.3287 \pm 0.0024$ | $0.2500 \pm 0.0024$ | $0.2049 \pm 0.0038$ | $0.2185 \pm 0.0056$ | $0.7622 \pm 0.0037$ |

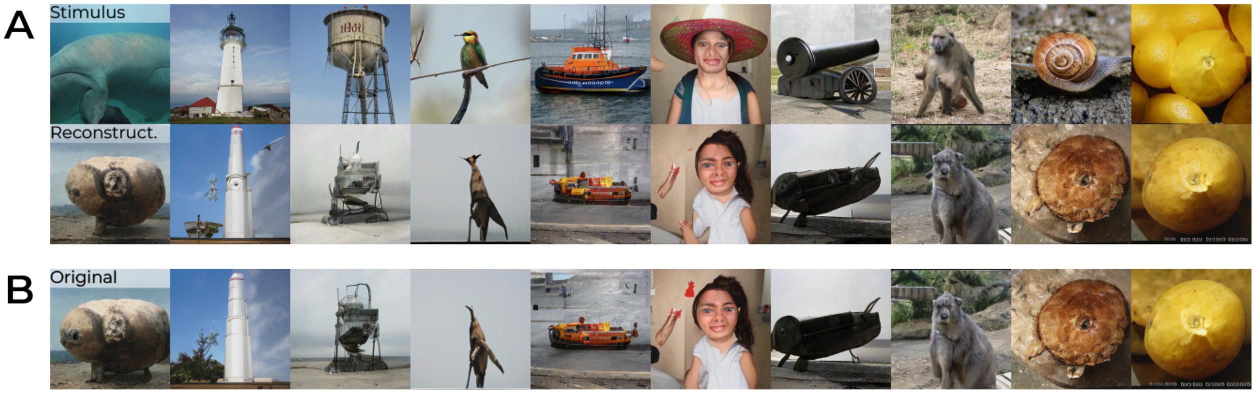


Figure 11: **S5.2 Qualitative reconstruction results:** test set stimuli (top) and their reconstructions from brain activity in V1, V4 and IT when the training examples of their class are excluded from training (middle). The original reconstructions, when all classes are included during training, are also displayed for reference.

#### S6 Permutation test analysis

VGG16 layer 1 cosine similarity
VGG16 layer 2 cosine similarity
VGG16 layer 3 cosine similarity
VGG16 layer 4 cosine similarity
VGG16 layer 5 cosine similarity
Latent cosine similarity
Random
Decoded from brain activity

*Legend*

| | VGG16 layer 1 | VGG16 layer 2 | VGG16 layer 3 | VGG16 layer 4 | VGG16 layer 5 | Latent |
|---|---|---|---|---|---|---|
| *all* | 0.79 | 0.77 | 0.59 | 0.62 | 0.59 | 0.55 |
| *V1* | 0.64 | 0.68 | 0.49 | 0.5 | 0.44 | 0.5 |
| *V4* | 0.63 | 0.67 | 0.49 | 0.5 | 0.42 | 0.5 |
| *IT* | 0.71 | 0.71 | 0.51 | 0.53 | 0.44 | 0.52 |
| ***rand.*** | **0.5** | **0.58** | **0.36** | **0.32** | **0.27** | **0.18** |

*(rand. averaged over 100 permutations)*

| | VGG16 layer 1 | VGG16 layer 2 | VGG16 layer 3 | VGG16 layer 4 | VGG16 layer 5 | Latent |
|---|---|---|---|---|---|---|
| *all* | 0.41 | 0.33 | 0.26 | 0.22 | 0.25 | 0.8 |
| *V1* | 0.39 | 0.31 | 0.22 | 0.15 | 0.14 | 0.73 |
| *V4* | 0.38 | 0.31 | 0.23 | 0.16 | 0.16 | 0.76 |
| *IT* | 0.38 | 0.31 | 0.23 | 0.18 | 0.17 | 0.77 |
| ***rand.*** | **0.28** | **0.25** | **0.18** | **0.12** | **0.08** | **0.04** |

*(rand. averaged over 100 permutations)*

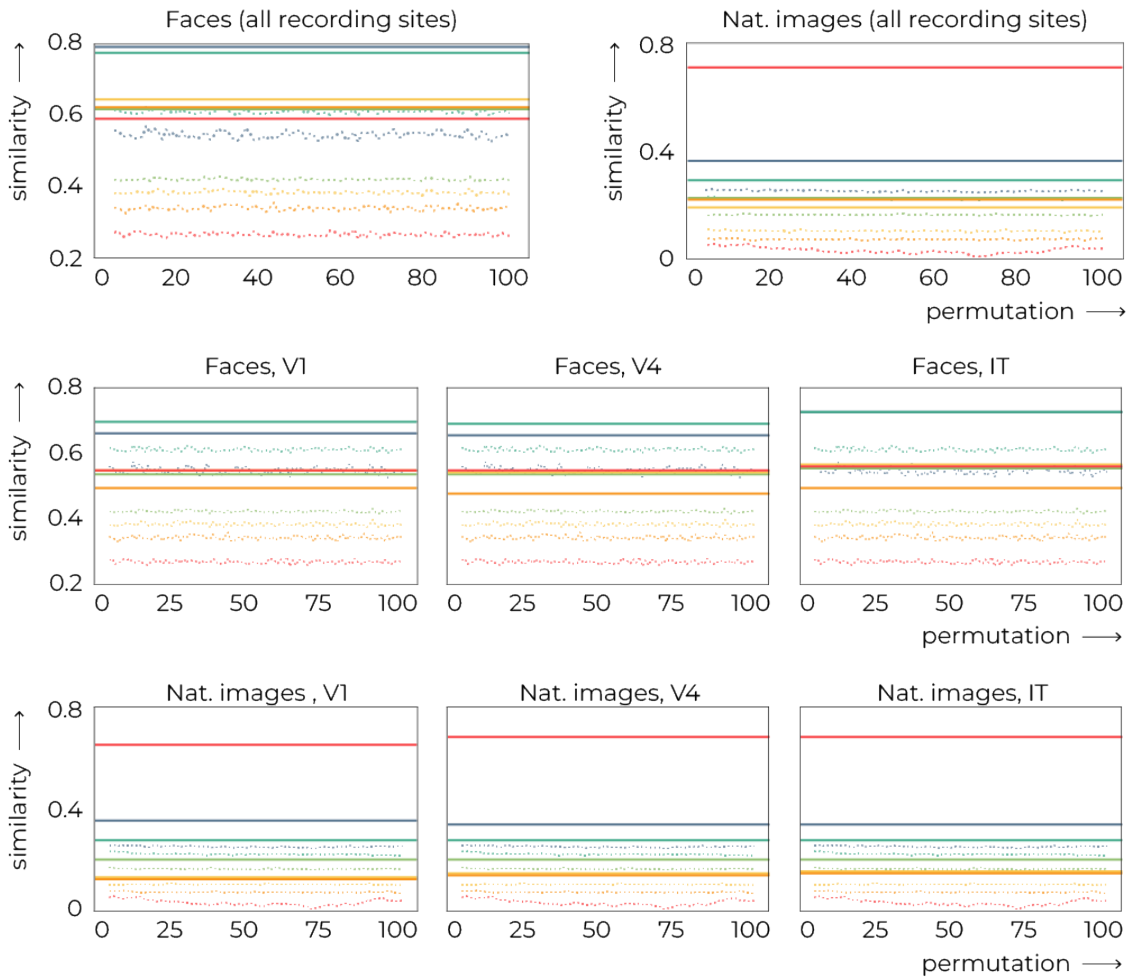


Figure 12: **S6.1 Permutation test analysis.** The quantitative results were verified with a permutation test as follows: per iteration, 100 and 200 latents (and their corresponding images) were randomly sampled for faces and natural images, respectively, to evaluate their similarity to the stimuli in terms of the six similarity metrics. In the above graphs, these similarity metrics were plotted over 100 iterations and we discovered that random samples were never better than our predictions from brain activity.

### S7 Visual guide

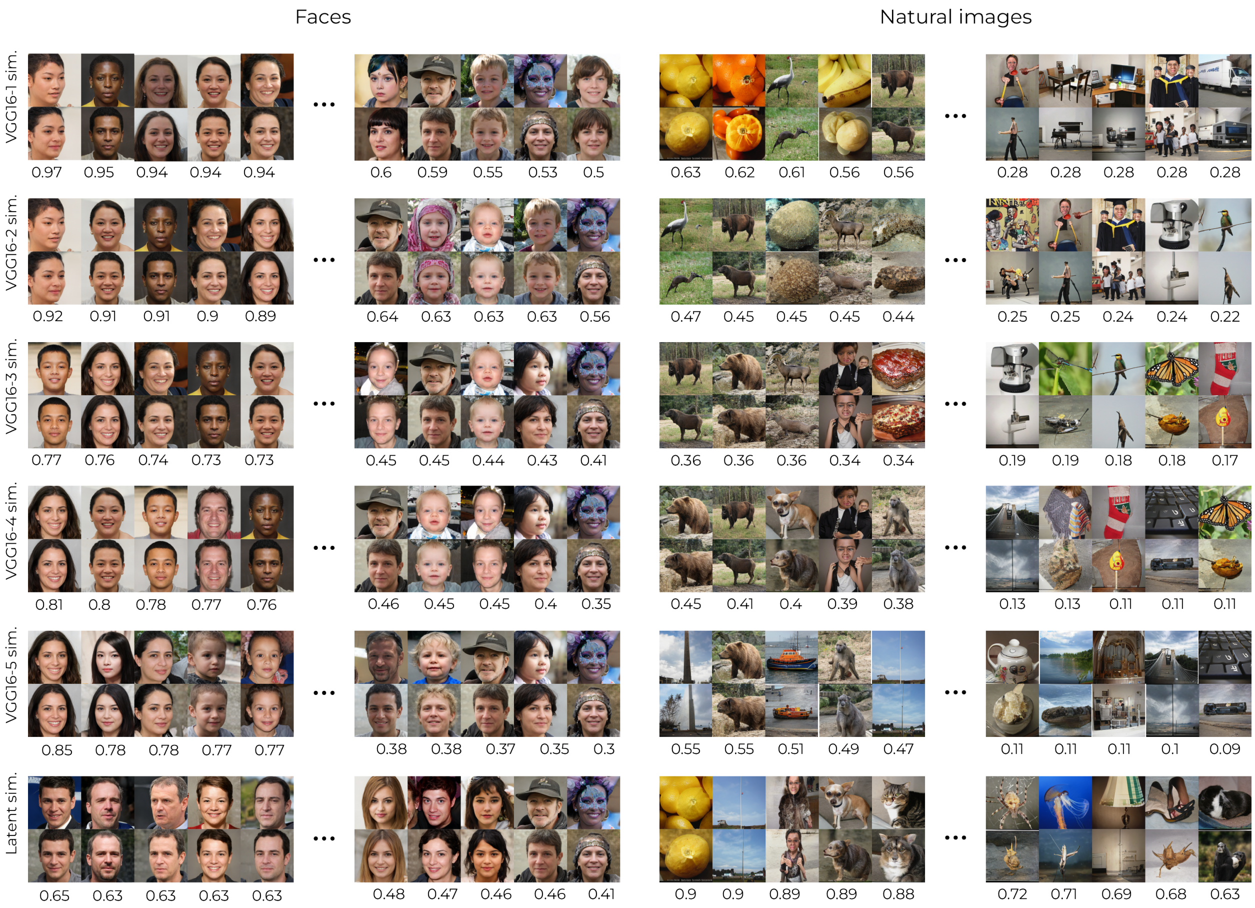


Figure 13: **7.1 Visual guide.** For the six similarity metrics, we display the 5 lowest and highest stimulus-reconstruction pairs from the dataset of faces- (left) and natural images dataset (right). The top row denotes the stimulus and the bottom row the reconstruction from brain activity.

#### S8 Abstract stimuli

Our experiment did not include abstract stimuli or optical illusions, as in [3]. It is however worth noting that the feasibility thereof, using StyleGAN-XL's framework, is an intriguing question. To this end, we (partially) leveraged the inversion script which can be found in the original GitHub repository of StyleGAN-XL. In brief, we optimized an input $w$-latent via the perceptual lpips loss using VGG16 features. We used the default parameters as specified in the script. Our findings suggest that, within the limitations of StyleGAN-XL's design tied to the natural image distribution, the generator indeed exhibits the capability to synthesize such images. This, in turn, offers an interesting perspective for future investigations in this direction. Note that inverting via the extended $W^+$ latent space could result in generated images that match the input images even more closely.

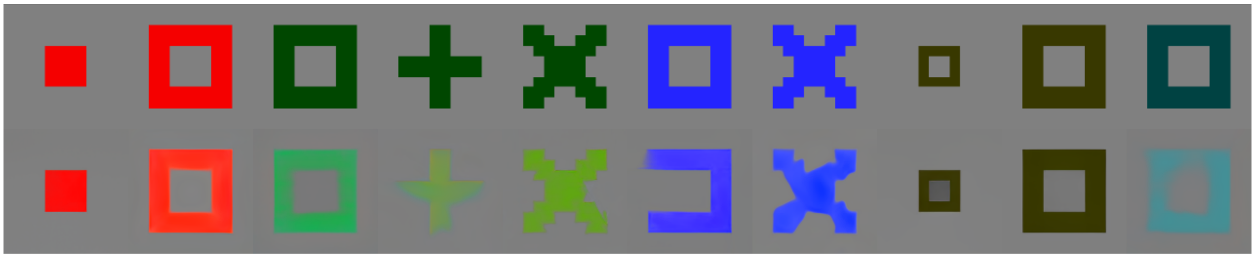

Figure 14: **S8.1 Generating abstract images.** Top: abstract image (taken from [3]). Bottom: image corresponding to the iteratively-optimized latent to match its visual features with those of the target latent.

#### S9 Category labels (Tiny ImageNet)

1. Egyptian cat
2. reel
3. volleyball
4. rocking chair
5. lemon
6. bullfrog
7. basketball
8. cliff
9. espresso
10. plunger
11. parking meter
12. German shepherd
13. dining table
14. monarch
15. brown bear
16. school bus
17. pizza
18. guinea pig
19. umbrella
20. organ
21. oboe
22. maypole
23. goldfish
24. potpie
25. hourglass
26. seashore
27. computer keyboard
28. Arabian camel
29. ice cream
30. nail
31. space heater
32. cardigan
33. baboon
34. snail
35. coral reef
36. albatross
37. spider web
38. sea cucumber
39. backpack
40. Labrador retriever
41. pretzel
42. king penguin
43. sulphur butterfly
44. tarantula
45. lesser panda
46. pop bottle
47. banana
48. sock
49. cockroach
50. projectile
51. beer bottle
52. mantis
53. freight car
54. guacamole
55. remote control
56. European fire salamander

57. lakeside
58. chimpanzee
59. pay-phone
60. fur coat
61. alp
62. lampshade
63. torch
64. abacus
65. moving van
66. barrel
67. tabby
68. goose
69. koala
70. bullet train
71. CD player
72. teapot
73. birdhouse
74. gazelle
75. academic gown
76. tractor
77. ladybug
78. miniskirt
79. golden retriever
80. triumphal arch
81. cannon
82. neck brace
83. sombrero
84. gasmask
85. candle
86. desk
87. frying pan
88. bee
89. dam
90. spiny lobster
91. police van
92. iPod
93. punching bag
94. beacon
95. jellyfish
96. wok
97. potter's wheel
98. sandal
99. pill bottle
100. butcher shop
101. slug
102. hog
103. cougar
104. crane
105. vestment
106. dragonfly
107. cash machine
108. mushroom
109. jinrikisha
110. water tower
111. chest
112. snorkel
113. sunglasses
114. fly
115. limousine
116. black stork
117. dugong

118. sports car
119. water jug
120. suspension bridge
121. ox
122. ice lolly
123. turnstile
124. Christmas stocking
125. broom
126. scorpion
127. wooden spoon
128. picket fence
129. rugby ball
130. sewing machine
131. steel arch bridge
132. Persian cat
133. refrigerator
134. barn
135. apron
136. Yorkshire terrier
137. swimming trunks
138. stopwatch
139. lawn mower
140. thatch
141. fountain
142. black widow
143. bikini
144. plate
145. teddy
146. barbershop
147. confectionery
148. beach wagon
149. scoreboard
150. orange
151. flagpole
152. American lobster
153. trolleybus
154. drumstick
155. dumbbell
156. brass
157. bow tie
158. convertible
159. bighorn
160. orangutan
161. American alligator
162. centipede
163. syringe
164. go-kart
165. brain coral
166. sea slug
167. cliff dwelling
168. mashed potato
169. viaduct
170. military uniform
171. pomegranate
172. chain
173. kimono
174. comic book
175. trilobite
176. bison
177. pole
178. boa constrictor

179. poncho
180. bathtub
181. grasshopper
182. walking stick
183. Chihuahua
184. tailed frog
185. lion
186. altar
187. obelisk
188. beaker
189. bell pepper
190. bannister
191. bucket
192. magnetic compass
193. meat loaf
194. gondola
195. standard poodle
196. acorn
197. lifeboat
198. binoculars
199. cauliflower
200. African elephant

#### References

[1] Shinji Nishimoto, An T Vu, Thomas Naselaris, Yuval Benjamini, Bin Yu, and Jack L Gallant. Reconstructing visual experiences from brain activity evoked by natural movies. *Current biology*, 21(19):1641–1646, 2011.

[2] Thomas Naselaris, Ryan J Prenger, Kendrick N Kay, Michael Oliver, and Jack L Gallant. Bayesian reconstruction of natural images from human brain activity. *Neuron*, 63(6):902–915, 2009.

[3] Guohua Shen, Tomoyasu Horikawa, Kei Majima, and Yukiyasu Kamitani. Deep image reconstruction from human brain activity. *PLoS computational biology*, 15(1):e1006633, 2019.
